# Genotypic complexity of Fisher’s geometric model

**DOI:** 10.1101/096438

**Authors:** Sungmin Hwang, Su-Chan Park, Joachim Krug

## Abstract

Fisher’s geometric model was originally introduced to argue that complex adaptations must occur in small steps because of pleiotropic constraints. When supplemented with the assumption of additivity of mutational effects on phenotypic traits, it provides a simple mechanism for the emergence of genotypic epistasis from the nonlinear mapping of phenotypes to fitness. Of particular interest is the occurrence of reciprocal sign epistasis, which is a necessary condition for multipeaked genotypic fitness landscapes. Here we compute the probability that a pair of randomly chosen mutations interacts sign-epistatically, which is found to decrease with increasing phenotypic dimension *n*, and varies non-monotonically with the distance from the phenotypic optimum. We then derive expressions for the mean number of fitness maxima in genotypic landscapes composed of all combinations of *L* random mutations. This number increases exponentially with *L*, and the corresponding growth rate is used as a measure of the complexity of the landscape. The dependence of the complexity on the model parameters is found to be surprisingly rich, and three distinct phases characterized by different landscape structures are identified. Our analysis shows that the phenotypic dimension, which is often referred to as phenotypic complexity, does not generally correlate with the complexity of fitness landscapes and that even organisms with a single phenotypic trait can have complex landscapes. Our results further inform the interpretation of experiments where the parameters of Fisher's model have been inferred from data, and help to elucidate which features of empirical fitness landscapes can be described by this model.

---

**A** fundamental question in the theory of evolutionary adaptation concerns the distribution of mutational effect sizes and the relative roles of mutations of small versus large effects in the adaptive process (Orr 2005). In his seminal 1930 monograph, Ronald Fisher devised a simple geometric model of adaptation in which an organism is described by *n* phenotypic traits and mutations are random displacements in the trait space (Fisher 1930). Each trait has a unique optimal value and the combination of these values defines a single phenotypic fitness optimum that constitutes the target of adaptation. Because random mutations act pleiotropically on multiple traits, the probability that a given mutation brings the phenotype closer to the target decreases with increasing *n*. Fisher’s analysis showed that, for large *n*, the mutational step size in units of the distance to the optimum must be smaller than 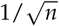 in order for the mutation to be beneficial with an appreciable probability. He thus concluded that the evolution of complex adaptations involving a large number of traits must rely on mutations of small effect. This conclusion was subsequently qualified by the realization that small effect mutations are likely to be lost by genetic drift, and therefore mutations of intermediate size contribute most effectively to adaptation (Kimura 1983; Orr 1998, 2000).

During the past decade Fisher's geometric model (FGM) has become a standard reference point for theoretical and experimental work on fundamental aspects of evolutionary adaptation (Tenaillon 2014). In particular, it has been found that FGM provides a versatile and conceptually simple mechanism for the emergence of epistatic interactions between genetic mutations in their effect on fitness (Martin *et al*. 2007; Gros *et al*.2009; Blanquart *et al*. 2014). For this purpose two extensions of Fisher's original formulation of the model have been suggested. First, phenotypes are assigned an explicit fitness value, which is usually taken to be a smooth function on the trait space with a single maximum at the optimal phenotype. Second, and more importantly, mutational effects on the phenotypes are assumed to be additive. As a consequence, any deviations from additivity that arise on the level of fitness are solely due to the nonlinear mapping from phenotype to fitness, or, in mathematical terms, due to the curvature of the fitness function. Because the curvature is largest around the phenotypic optimum, epistasis generally increases upon approaching the optimal phenotype and is weak far away from the optimum. Several recent studies have made use of the framework of FGM to interpret experimental results on pairwise epistastic interactions and to estimate the parameters of the model from data (Martin *et al*. 2007; Velenich and Gore 2013; Weinreich and Knies 2013; Perfeito *et al*. 2014; Schoustra *et al*. 2016).

A particularly important form of epistatic interaction is sign epistasis, where a given mutation is beneficial or deleterious depending on the genetic background (Weinreich *et al*. 2005). Two types of sign epistasis are distinguished depending on whether one of the mutations affects the effect sign of the other but the reverse is not true (*simple sign epistasis*), or whether the interaction is reciprocal (*reciprocal sign epistasis*); for a pictorial representation of the two kinds of sign epistasis, see, for example, Poelwijk *et al*. (2007). Sign epistasis can arise in FGM either between large effect beneficial mutations that in combination overshoot the fitness optimum, or between mutations of small fitness effect that display antagonistic pleiotropy (Blanquart *et al*.2014). The presence of sign epistasis is a defining feature of genotypic fitness landscapes that are complex, in the sense that not all mutational pathways are accessible through simple hill-climbing and multiple genotypic fitness peaks may exist (Weinreich *et al*. 2005; Franke *et al*. 2011; de Visser and Krug 2014). Specifically, reciprocal sign epistasis is a necessary condition for the existence of multiple fitness peaks (Poelwijk *et al*.2011; Crona *et al*. 2013).

Following a common practice, here a genotypic fitness landscape is understood to consist in the assignment of fitness values to all combinations of *L* haploid, biallelic loci that together constitute the *L*-dimensional genotype space. A peak in such a landscape is a genotype that has higher fitness than all its *L* neighbors that can be reached by a single point mutation (Kauffman and Levin 1987). Note that, in contrast to the continuous phenotypic space on which FGM is defined, the space of genotypes is discrete.

Blanquart *et al*. (2014) showed that an ensemble of *L*-dimensional genotypic landscapes can be constructed from FGM by combining subsets of *L* randomly chosen mutational displacements. Each sample of *L* mutations defines another realization of the landscape ensemble, and the exploratory simulations reported by Blanquart *et al*. (2014) indicate a large variability among the realized landscapes. Nevertheless some general trends in the properties of the genotypic landscapes were identified. In particular, as expected on the basis of the considerations outlined above, the genotypic landscapes are essentially additive when the focal phenotype representing the unmutated wild type is far away from the optimum and become increasingly rugged as the optimal phenotype is approached.

In this article we present a detailed and largely analytic study of the properties of genotypic landscapes generated under FGM. The focus is on two types of measures of landscape complexity, that is, the fraction of sign-epistatic pairs of random mutations and the number of fitness maxima in the genotypic landscape. A central motivation for our investigation is to assess the potential of FGM and related phenotypic models to explain the properties of empirical genotypic fitness landscapes of the kind that have been recently reported in the literature (Szendro *et al*. 2013; Weinreich *et al*. 2013; de Visser and Krug 2014). The ability of nonlinear phenotype-fitness maps to explain epistatic interactions among multiple loci has been demonstrated for a virus (Rokyta *et al*. 2011) and for an antibiotic resistance enzyme (Schenk *et al*. 2013), but a comparative study of several different data sets using Approximate Bayesian Computation has questioned the broader applicability of phenotype-based models (Blanquart and Bataillon 2016). It is thus important to develop a better understanding of the structure of genotypic landscapes generated by phenotypic models such as FGM.

In the next section we describe the mathematical setting and introduce the relevant model parameters: the phenotypic and genotypic dimensionalities *n* and *L*, the distance of the focal phenotype to the optimum, and the standard deviation of mutational displacements. As in previous studies of FGM, specific scaling relations among these parameters have to be imposed in order to arrive at meaningful results for large *n* and *L*. We then present analytic results for the probability of sign epistasis and the behavior of the number of fitness maxima for large *L*, both in the case of fixed phenotypic dimension *n* and for a situation where the joint limit *n, L* → ∞ is taken at constant ratio *α* = *n/L*.

Similar to other probabilistic models of genotypic fitness landscapes (Kauffman and Levin 1987; Weinberger 1991; Evans and Steinsaltz 2002; Durrett and Limic 2003; Limic and Pemantle 2004; Neidhart *et al*. 2014), the number of maxima generally increases exponentially with *L*, and we use the exponential growth rate as a measure of genotypic complexity. We find that this quantity displays several phase transitions as a function of the parameters of FGM which separate parameter regimes characterized by qualitatively different landscape structures. Depending on the regime, the genotypic landscapes induced by FGM become more or less rugged with increasing phenotypic dimension. This indicates that the role of the number of phenotypic traits in shaping the fitness landscapes of FGM is much more subtle than has been previously appreciated, and that the sweeping designation of *n* as (phenotypic) “complexity” can be misleading. Further implications of our study for the theory of adaptation and the interpretation of empirical data will be elaborated in **Discussion**.

## Model

### Basic properties of FGM

In FGM, the phenotype of an organism is modeled as a set of *n* real-valued traits and represented by a vector 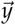 = (*y*_1_,*y*_2_,…,*y_n_*) in the *n* dimensional Cartesian space, 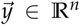. The fitness W 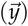 is assumed to be a smooth, single-peaked function of the phenotype 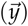. By choosing an appropriate coordinate system, the optimum phenotype, i.e., the combination of phenotypic traits with the highest fitness value, can be placed at the origin in 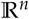. We also assume that the fitness *W* 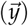 depends on the distance to the optimum 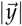 but not on the direction of 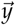, which can be justified by arguments based on random matrix theory (Martin 2014). The uniqueness of the phenotypic optimum at the origin implies that *W* 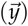 is a decreasing function of 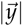. The form of the fitness function will be specified below when needed. Most of the results presented in this paper are however independent of the explicit shape of *W*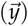, as they rely solely on the relative ordering of different genotypes with respect to their fitness.

When a mutation arises the phenotype of the mutant becomes 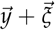, where 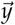 is the parental phenotype and the mutational vector 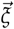 corresponds to the change of traits due to the mutation. The key result derived by Fisher (1930) concerns the fraction *P_b_* of beneficial mutations arising from a wild type phenotype located at distance d from the optimum. Assuming that mutational displacements have a fixed length 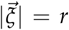 and random directions, he showed that for *n* ≫ 1

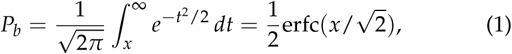
where erfc denotes the complementary error function and *x* = 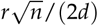. Thus, for large *n* the mutational step size has to be much smaller than the distance to the optimum, 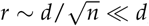, for the mutation to have a chance of increasing fitness.

As has become customary in the field, we here assume that the mutational displacements are independent and identically distributed (i.i.d.) random variables drawn from a *n*- dimensional Gaussian distribution with zero mean. The covariance matrix can be taken to be of diagonal form *σ*^2^*I*, where *I* is the *n*-dimensional identity matrix and *σ*^2^ is the variance of a single trait (Blanquart *et al*. 2014). In the limit *n* → ∞ the form of the distribution of the mutational displacements becomes irrelevant owing to the central limit theorem, and therefore Fisher's result Equation 1 holds also in the present setting of Gaussian mutational displacements of mean size 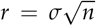(Waxman and Welch 2005; Ram and Hadany 2015); an explicit derivation will be provided below. Because lengths in the phenotype space can be naturally measured in units of *σ*, the parameters *d* and *σ* should always appear as the ratio *d/σ* as can be seen in Equation 1. Thus, without loss of generality, we can set *σ* = 1. In the following we denote the (scaled) wild type phenotype by 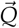, its distance to the optimum by

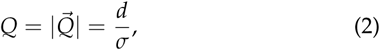
and draw the displacement vectors 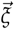 from the *n*-dimensional Gaussian 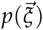 with unit covariance matrix.

By normalizing phenotypic distances to the standard deviation *σ* of the mutational effect on a single trait, we are adopting a particular pleiotropic scaling that has been referred to as the “Euclidean superposition model” (Wagner *et al*. 2008; Hermisson and McGregor 2008). An alternative choice which is closer to Fisher’s original formulation but appears to have less empirical support is the “total effect model”, where in the total length *r* of the mutational displacements is taken to be independent of *n*. Since 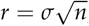 this implies that the single trait effect size decreases with *n* as 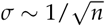 As a consequence the parameter Q defined by Equation 2 becomes *n*-dependent and increases as 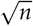 provided *d* does not depend on *n* (Orr 2000). The results presented below will always be given in terms of ratios of the basic parameters of FGM, such that their translation to the total effect model is in principle straightforward. We will nevertheless explicitly point out instances where the two settings give rise to qualitatively different behaviors.

**Figure 1.**
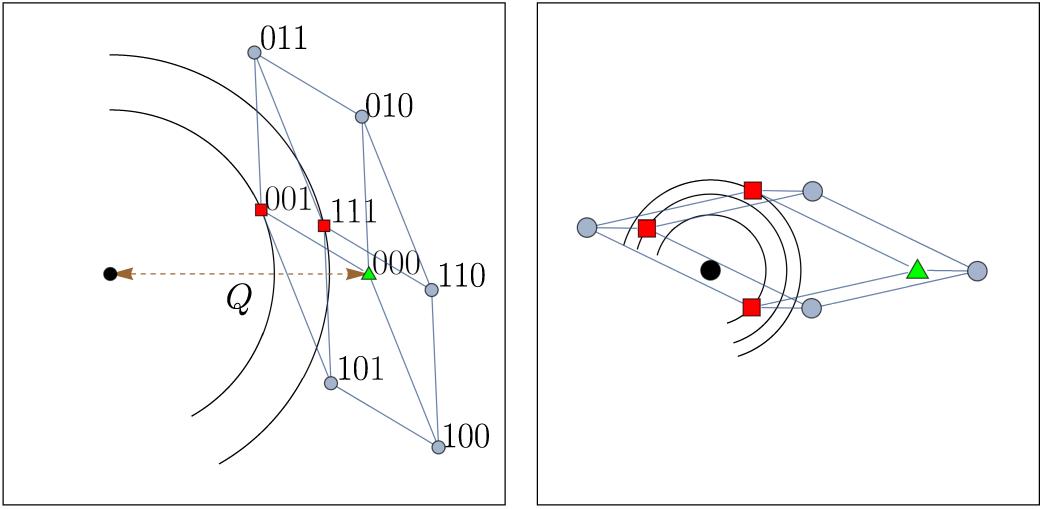
Examples of three-dimensional genotypic fitness landscapes induced by FGM with two phenotypic dimensions (*L* = 3 and *n* = 2). The panels show the projection of the discrete genotype space onto the phenotype plane, where the phenotypic optimum is represented by a black dot. In the left panel the binary sequence notation for genotypes is indicated. The wild type genotype 000, marked by a green triangle, is located at distance *Q* from the phenotypic optimum. The nodes represented by red squares are local fitness maxima of the genotypic landscapes, as can be seen from the contour lines of constant fitness. In the right panel the mutant phenotypes overshoot the optimum, whereas in the left panel they do not.

### The genotypic fitness landscape induced by FGM

In order to study epistasis within FGM, Fisher’s original definition has to be supplemented with a rule for how the effects of multiple mutations are combined. Based on earlier work (Lande 1980) in quantitative genetics, Martin *et al*. (2007) introduced the assumption that mutations act additively on the level of the phenotype. Thus the phenotype arising from two mutations 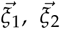 applied to the wild type 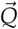 is simply given by 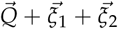. This definition suffices to associate an *L*-dimensional genotypic fitness landscape to any set of *L* mutational displacements 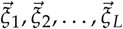 (Blanquart *et al*. 2014). For this purpose the haploid genotype *τ* is represented by a binary sequence with length *L, τ***=** (*τ*_1_, *τ*_2_,…, *τ_L_*) with τ*_i_* = 1 (τ*_i_* = 0) in the presence (absence) of the *i*'th mutation. For the wild type τ*_i_* = 0 for all *i*, and in general the phenotype vector associated with the genotype *τ* reads 
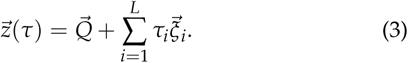

Two examples illustrating this genotype-phenotype map and the resulting genotypic fitness landscapes with *L* = 3 and *n* = 2 are shown in Figure 1.

Since fitness decreases monotonically with the distance to the optimum phenotype, a natural proxy for fitness is the negative squared magnitude of the phenotype vector

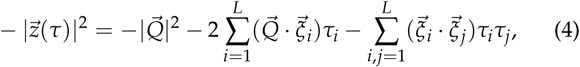
where 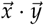 denotes the scalar product between two vectors 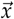 and 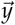. This quantity is thus seen to consist of a part that is additive across loci with coefficients given by the scalar products 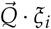 and a pairwise epistatic part with coefficients 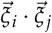.

It is instructive to decompose Equation 4 into contributions from the mutational displacements parallel and perpendicular to 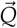. Writing 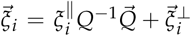 with 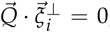 Equation 4 can be recast into the form

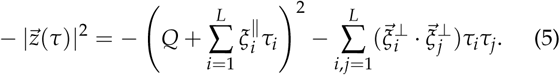

The first term on the right hand side contains both additive and epistatic contributions associated with displacements along the 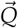-direction. The second term is dominated by the diagonal contributions with *i***=***j* and is of order *L*(*n*— 1) because 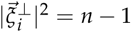 on average.

We now show how the first term on the right hand side of Equation 5 can be made to vanish for a range of *Q*. For this purpose consider the subset of phenotypic displacement vectors for which the component 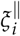 in the direction of 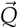 is negative. There are on average *L*/2 such mutations, and the expected value of each component is

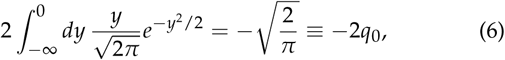
where the factor 2 in front of the integral arises from conditioning on 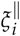 < 0. Setting *τ_i_***=** 1 for ***s*** out of these *L*/2 vectors and *τ_i_***=** 0 for all other mutations, the sum inside the brackets in Equation 5 becomes approximately equal to —2*q*_0_s which cancels the *Q*-term for *s***=** *Q*/2q_0_. Since *s* can be at most *L*/2 in a typical realization, such genotypes can be constructed with a probability approaching unity provided *Q***<***q*_0_*L*.

We will see below that the structure of the genotypic fitness landscapes induced by FGM depends crucially on whether or not the phenotypes of multiple mutants are able to closely approach the phenotypic optimum. Assuming that the contributions from the perpendicular displacements in Equation 5 can be neglected, which will be justified shortly, the simple argument given above shows that a close approach to the optimum is facile when *Q***<***q*_0_*L*. but becomes unlikely when *Q*≫*q*_0_*L*.This observation hints at a possible transition between different types of landscape topographies at some value of *Q* which is proportional to *L*. The existence and nature of this transition is a central theme of this paper.

### Scaling limits

Since we are interested in describing complex organisms with large phenotypic and genotypic dimensions, appropriate scaling relations have to be imposed to arrive at meaningful asymptotic results. Three distinct scaling limits will be considered.

(i) Fisher’s classic result (Equation 1) shows that the distance of the wild type from the phenotypic optimum has to be increased with increasing *n* in order to maintain a nonzero fraction of beneficial mutations for *n*→ ∞ to. In our notation Fisher’s parameter is

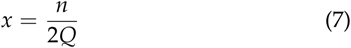
and hence *Fisher scaling* implies taking *n, Q*→ ∞ to at fixed ratio *n*/*Q*. We will extend Fisher’s analysis by computing the probability of sign epistasis between pairs of mutations for fixed *x* and large *n*, which amounts to characterizing the shape of genotypic fitness landscapes of size *L* = 2.

(ii) We have argued above that the distance towards the phenotypic optimum that can be covered by typical multiple mutations is of order *L*, and hence the limit *L →*∞ is naturally accompanied by a limit *Q* → ∞ at fixed ratio

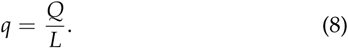

From a biological point of view one expects that *L* ≫ *n* ≫ 1, which motivates to consider the limit *L, Q→ ∞* at constant phenotypic dimension *n*. Under this scaling the first term on the right hand side of Equation 5 is of order *L*^2^, whereas the contribution from the perpendicular displacements is only (*n*— 1)*L*. Thus in this regime the topography of the fitness landscape is determined mainly by the one-dimensional mutational displacements in the 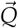-direction, which is reflected by the fact that the genotypic complexity is independent of *n* to leading order and coincides with its value for the case *n***=** 1, in which the perpendicular contribution in Equation 5 does not exist (see **Results**).

(iii) By contrast, the perpendicular displacements play an important role when both the phenotypic and genotypic dimensions are taken to infinity at fixed ratio 
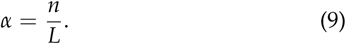

Combining this with the limit *Q → ∞* at fixed *q* = *Q/L*, both terms on the right hand side of Equation 5 are of the same order ∼ *L*^2^. Fisher’s parameter (Equation 7) is then also a constant given by *x= α* / (2*q*).

### Preliminary considerations about genotypic fitness maxima

To set the stage for the detailed investigation of the number of genotypic fitness maxima in **Results**, it is useful to develop some intuition for the behavior of this quantity based on the elementary properties of FGM that have been described so far. For this purpose we consider the probability *P*_wt_ for the wild type to be a local fitness maximum, which is equal to the probability that all the *L* mutations are deleterious. Since mutations are statistically independent, we have 
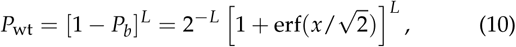
where erf = 1 – erfc is the error function. Under the (highly questionable) assumption that this estimate can be applied to all 2*^L^* genotypes in the landscape, we arrive at the expression

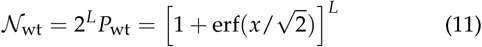
for the expected number of genotypic fitness maxima.

Consider first the scaling limit (ii), where *x* **=** *n***/**(2*Q*) = *n/*(*2qL*)→ 0. Expanding the error function for small arguments as 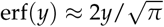 we obtain

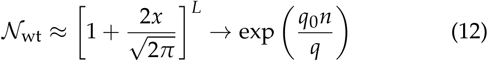
for *L*→∞, where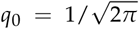 was defined in Equation 6. We will show below that this expression correctly captures the asymptotic behavior for very large *q* but generally grossly underestimates the number of maxima. The reason is that for moderate values of *q* (in particular for *q* < *q*_0_) the relevant mutant phenotypes are much closer to the origin than the wild type, which entails a mechanism for generating a large number of fitness maxima that grows exponentially with *L*.

Such an exponential dependence on *L* is expected from Equation 11 in the scaling limit (iii), where *x = α****/*** (2*q*) is a nonzero constant and the expression in the square brackets is larger than 1. Although this general prediction is confirmed by the detailed analysis for this case, the behavior of the number of maxima predicted by Equation 11 will again turn out to be valid only when *q* is very large. In particular, whereas Equation 11 is an increasing function of *α* for any *q*, we will see below that the expected number of maxima actually decreases with increasing phenotypic dimension (hence increasing *α*) in a substantial range of *q*. In qualitative terms, this can be attributed to the effect of the perpendicular displacements in Equation 5, which grows with *α* and makes it increasingly more difficult for the mutant phenotypes to closely approach the origin.

**Table 1.**
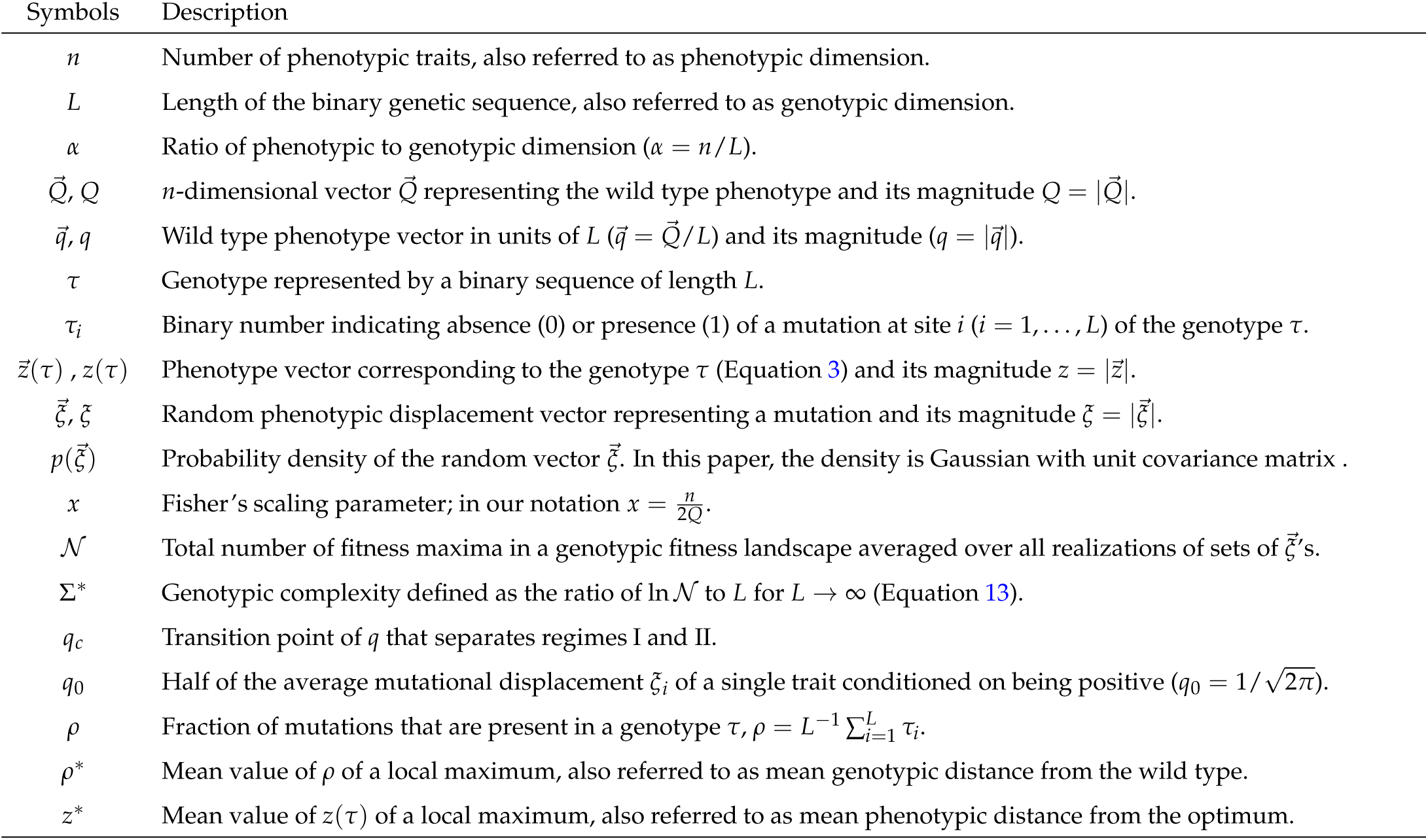
List of Mathematical Symbols

The observation that the number of genotypic fitness maxima grows exponentially with *L* in most cases motivates to make use of the corresponding growth rate as a measure of the ruggedness of the landscape. We therefore define the *genotypic complexity* Σ* through the limiting relation

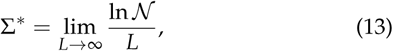
where 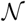 is the *average* number of genotypic fitness maxima and *L* is the sequence length. Since the total number of binary genotypes is 2*^L^*, the complexity is bounded from above by ln 2. If any genotype had the same probability *P*_max_ of being a fitness maximum (which is in fact not the case for FGM), we could write 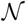 and hence *P*_max_ ˜ exp[—(ln2 — Σ*)*L*].

### Data availability

The authors state that all data necessary for confirming the conclusions presented in the article are represented fully within the article. All numerical calculations including simulations^1^ described in this work were implemented in Mathematica and C++. All relevant source codes are available upon request.

## Results

### Preliminary note

In the following sections our results on the structure of genotypic fitness landscapes induced by FGM are stated in precise mathematical terms and the key steps of their derivation are outlined, with some technical details relegated to the Appendices. In order to facilitate the navigation through the inevitable mathematical formalism, we display the definitions of the most commonly used mathematical symbols in Table 1. Moreover, we provide numbered summaries at the end of each subsection which state the main results without resorting to mathematical expressions.

### Sign epistasis

We first study the local topography of the fitness landscape around the wild type, focusing on the epistasis between two random mutations with phenotypic displacements 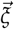 and 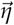. Since fitness is determined by the magnitude of a phenotypic vector, i.e., the distance of the phenotype from the origin, the epistatic effect of the two mutations can be understood by analyzing how the magnitudes of the four vectors 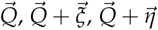 and 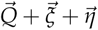 are ordered. To this end, we introduce the quantities

**Figure 2.**
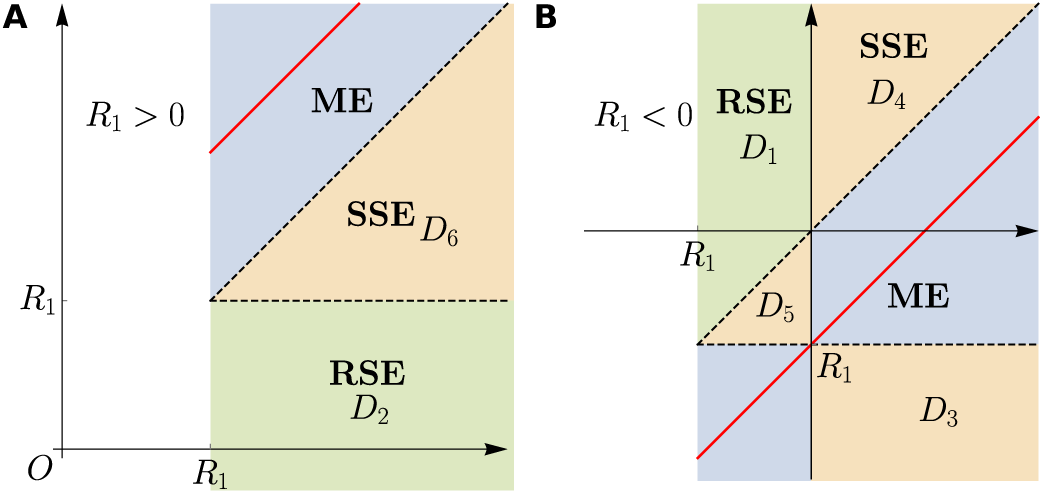
Domains in the (*R*_2_, *R*)-plane contributing to different types of epistasis: magnitude epistasis (ME), simple sign epistasis (SSE) and reciprocal sign epistasis (RSE). The two panels illustrate the two cases: (**A**) R_1_ > 0 and (**B**) R_1_ < 0. The red solid lines indicate *R***=***R_1_* + R_2_. The labeling of the domains *D*_1_, …, *D*_6_ is used in the derivation in Appendix B.

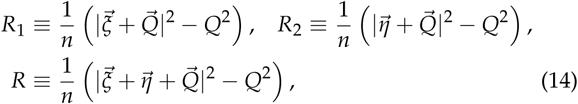
where division by *n* guarantees the existence of a finite limit for *n →* ∞. The sign of these quantities determines whether a mutation is beneficial or deleterious. For example, if *R*_1_ < 0, the mutation 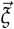 is beneficial; if *R* **>** 0 the two mutations combined together confer a deleterious effect; and so on. We will see later that R_1,2_ and *R* are actually closely related to the selection coefficients of the respective mutations.

We proceed to express the different types of pairwise epis-tasis defined by Weinreich *et al*. (2005) and Poelwijk *et al*. (2007) in terms of conditions on the quantities defined in Equation 14. Without loss of generality we assume *R*_1_ < *R*_2_ and consider first the case where both mutations are beneficial, *R*_1_ < *R*_2_ < 0. Then magnitude epistasis (ME), the absence of sign epistasis, applies when the fitness of the double mutant is higher than that of each of the single mutants, i.e., *R* < *R*_1_ < *R*_2_ < 0. Similarly for two deleterious mutations the condition for ME reads *R* > *R*_2_ > *R*_1_ > 0. When one mutant is deleterious and the other beneficial, in the case of ME the double mutant fitness has to be intermediate between the two single mutants, which implies that *R_1_* < *R* **<** *R_2_* when *R*_2_ > 0 > *R_1_*.

The condition for reciprocal sign epistasis (RSE) reads *R* > *R*_2_ > *R*_1_ when both single mutants are beneficial and *R* < *R*_1_ < *R*_2_ when both are deleterious, and the remaining possibility *R*_1_ < *R* < *R*_2_ corresponds to simple sign epistasis (SSE) between two mutations of the same sign. If the two single mutant effects are of different signs, RSE is impossible and SSE applies when *R* **<** *R_1_* **<** 0 < *R*_2_ or *R* **>** *R_2_* **>** 0 > *R*_1_. Figure 2 depicts the different categories of epistasis as regions in the (*R*_2_, *R*)-plane. Note that the corresponding picture for *R*_1_ > *R*_2_ is obtained by exchanging *R*_1_ ↔ *R*_2_.

To find the probability of each epistasis, we require the joint probability density 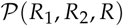. In Appendix A it is shown

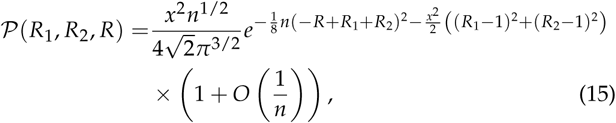
which can be obtained rather easily by resorting to the central limit theorem (CLT). The applicability of the CLT follows from the fact that *R*_1,2_ and *R* are sums of a large number of independent terms for *n*→*∞* (Waxman and Welch 2005; Ram and Hadany 2015). According to the CLT, it is sufficient to determine the first and second cumulants of these quantities. Denoting averages by angular brackets, we find the mean 〈*R_i_*〉 = 1, the variance 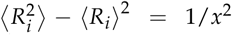, and the covariance 〈*R*_1_*R*_2_ 〉 – 〈*R*_1_〉 〈R_2_〉= 0 (*i* = 1,2). Similarly, the corresponding quantitites evaluated for *R* – *R*_1_ – *R*_2_ are 〈*R* – *R*_1_ – *R*_2_〉 = 0, 〈(*R* – *R*_1_ – *R*_2_)^2^〉 – 〈*R* – *R*_1_ – *R*_2_〉^2^ = 4/*n*, and 〈(*R* – *R*_1_ – *R_2_*)*R_i_*〉 – 〈*R* – *R_1_* – *R_2_*〉 〈*R_i_*〉= 0 (*i* = 1,2). With an appropriate normalization constant, this leads directly to Equation 15.

As a first application, we rederive Fisher’s Equation 1 by integrating 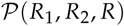 over the region *R*_1_ < 0 for all *R*_2_ and *R*, which indeed yields

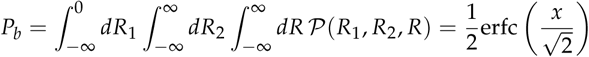

An immediate conclusion from the form of 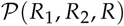 is that it is unlikely to observe sign epistasis for large *n*, because 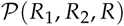 becomes concentrated along the line *R***=***R_1_* + *R*_2_ as *n* increases. As can be seen in Figure 2, this line touches the region of SSE in one point for *R*_1_ < 0, whereas it maintains a finite distance to the region of RSE everywhere. This indicates that the probability of reciprocal sign epistasis decays more rapidly with increasing *n* than the probability of simple sign epistasis. Moreover, one expects the latter probability to be proportional to the width of the region around the line *R***=***R_1_* + *R*_2_ where the joint probability in Equation 15 has appreciable weight, which is of order 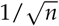

To be more quantitative, we need to integrate 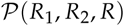 over the domains in Figure 2 corresponding to the different categories of epistasis. In Appendix B, we obtain the asymptotic expressions

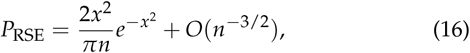

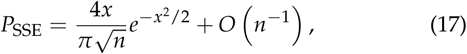
for the probabilities of reciprocal (*P*_RSE_) and simple (*P*_SSE_) sign epistasis. Due to the non-linearity of the phenotype-fitness map, FGM does not allow for strictly non-epistatic combination of fitness effects. The probability of magnitude epistasis, therefore, is given by *P*_ME_ = 1 − *P*_RSE_ − *P*_SSE_. Interestingly, the probability of sign epistasis varies non-monotonically with *x*. To confirm our analytic results, we compare our results with simulations in Figure 3, which shows an excellent agreement.

Similarly, we can calculate the probabilities of sign epistasis conditioned on both mutations being beneficial, which in our setting means *R*_2_ < 0. The conditioning requires normalization by the unconditional probability of two random mutations being beneficial, which is given by the square of *P_b_* in Equation 1.

**Figure 3.**
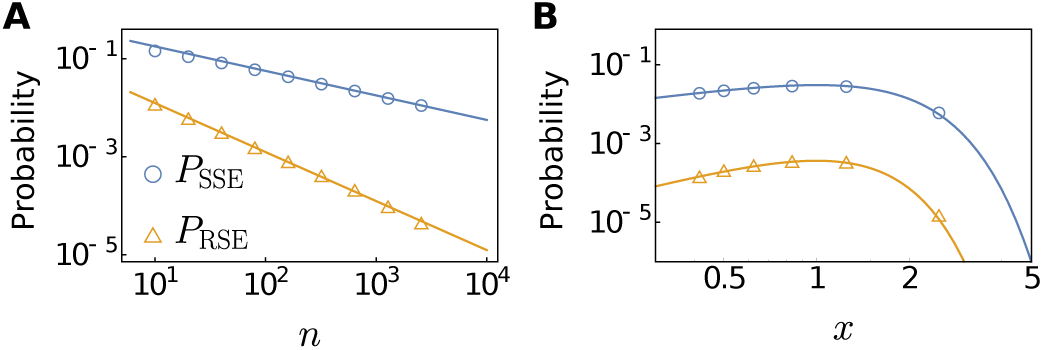
Comparison of analytic results for the probability of epistasis with simulations. Depicted are probabilities of simple (*P*_SSE_) and reciprocal (*P*_RSE_) sign epistasis between two randomly chosen mutations among nearest neighbor genotypes of the wild type (**A**) as functions of *n* for fixed Fisher parameter *x* = 0.5 and (**B**) as function of *x* for fixed phenotypic dimension *n* = 640. For each parameter set, 10^4^ randomly generated landscapes were analyzed. The asymptotic expressions provide accurate approximations even for moderate *n* > 10. The non-monotonic behavior with respect to *x* means that the probabilities are non-monotonic functions of *Q* for fixed n and vice versa.

Hence

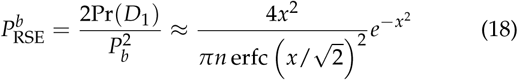
And 
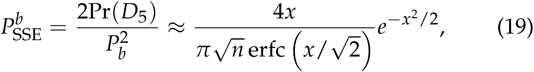
where Pr(*D_i_*) denotes the integral of the joint probability density over the domain *D_i_* in Figure 2 (see Appendix B).

As anticipated from the form of Equation 15, the fraction of sign epistatic pairs of mutations decreases with increasing phenotypic dimension *n*, and this decay is faster for reciprocal sign epistasis (∼ 1/*n*) than for simple sign epistasis 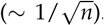 At first glance this might seem to suggest that FGM has little potential for generating rugged genotypic fitness landscapes. However, as we will see below, the results obtained in this section apply only to the immediate neighborhood of the wild type phenotype. They are modified qualitatively in the presence of a large number of mutations that are able to substantially displace the phenotype and allow it to approach the phenotypic optimum.

As a slight variation to the previous setting, one may consider the fraction of sign epistasis conditioned on the two single mutations to have the same selection strength, as recently investigated by Schoustra *et al*. (2016). In our notation this implies that 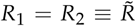, and it is easy to see that sign epistasis is always reciprocal in this case. If the two mutations are beneficial 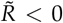, and the condition for (reciprocal) sign epistasis is 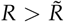 The corresponding probability is

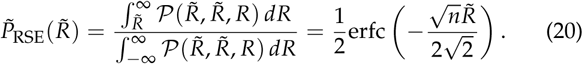

Following the same procedure for deleterious mutations 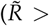 one finds that the probability is actually symmetric around 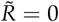 and hence depends only on 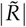.

In order to express 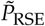 in terms of the selection coefficient of the single mutations we introduce a Gaussian phenotypic fitness function of the form

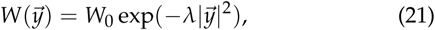
where λ > 0 is a measure for the strength of selection. The selection coefficient of a mutation with phenotypic effect 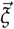 is then given by 
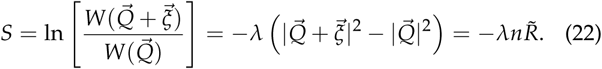

**Figure 4.**
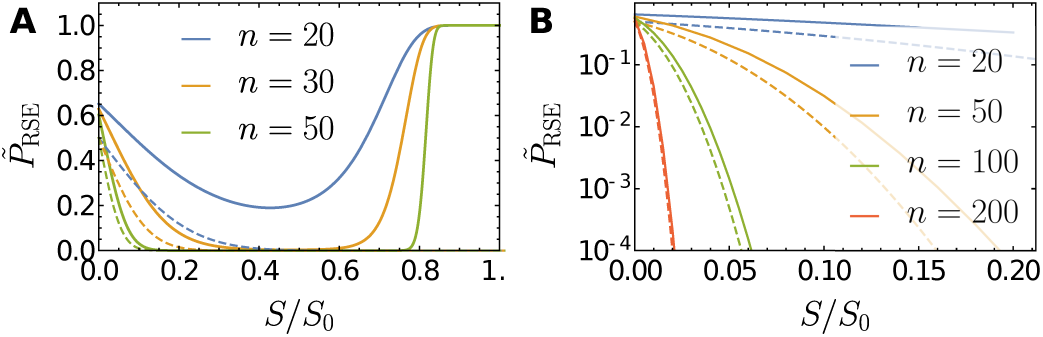
Probability of reciprocal sign epistasis 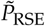 conditioned on the selection coefficients S of the two single mutations to be equal and positive. Here, the fitness of a phenotype 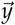 is assumed to be 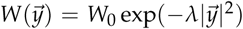, where the parameter λ is related to the maximal beneficial selection coefficient *S*_0_ through the relation *S*_0_ = *λQ*^2^. Dashed lines depict the asymptotic expression Equation 23, and solid lines were obtained numerically using the Gaussian approximation for the distribution of epistasis developed by Schoustra *et al*. (2016).

To fix the value of λ we note that the largest possible selection coefficient, which is achieved for mutations that reach the phenotypic optimum, is *S*_0_ = λQ^2^, and hence 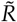 is related to the selection coefficient through 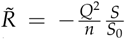 With this substitution the result in Equation 20 becomes

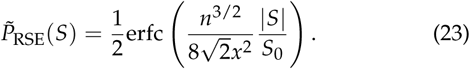

The probability of sign epistasis conditioned on selection strength takes on its maximal value 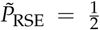 in the neutral limit *S* → 0 and decreases monotonically with |*S*|. Similar to the results of Equation 16, Equation 17 and Equation 18 for unconstrained mutations, it also decreases with increasing phenotypic dimension *n* when *S* and *x* are kept fixed.

In a previous numerical study carried out at finite *Q* and *n* it was found that 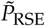 varies non-monotonically with *S* for the case of beneficial mutations, and displays a second peak at the maximum selection coefficient *S* = *S*_0_ (Schoustra *et al*. 2016). The two peaks were argued to reflect the two distinct mechanisms giving rise to sign epistasis within FGM (Blanquart *et al*. 2014). Mutations of small effect correspond to phenotypic displacements that proceed almost perpendicular to the direction of the phenotypic optimum, and sign epistasis is generated through antagonistic pleiotropy. On the other hand, for mutations of large effect the dominant mechanism for sign epistasis is through overshooting of the phenotypic optimum. Because of the Fisher scaling implemented in this section with *Q, n* → ∞ at fixed *x* = *n*/(2*Q*), the second class of mutations cannot be captured by our approach and only the peak at small *S* remains. Figure 4 (A) shows the full two-peak structure for a few representative values of *n*, and Figure 4 (B) illustrates the convergence to the asymptotic expression Equation 23 for the left peak. Using the results of Schoustra *et al*. (2016), it can be shown that the right peak becomes a step function for *n* → ∞, displaying a discontinuous jump from 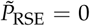 to 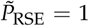 at *S*/*S*_0_ = 8/9 = 0.888….

**Figure 5.**
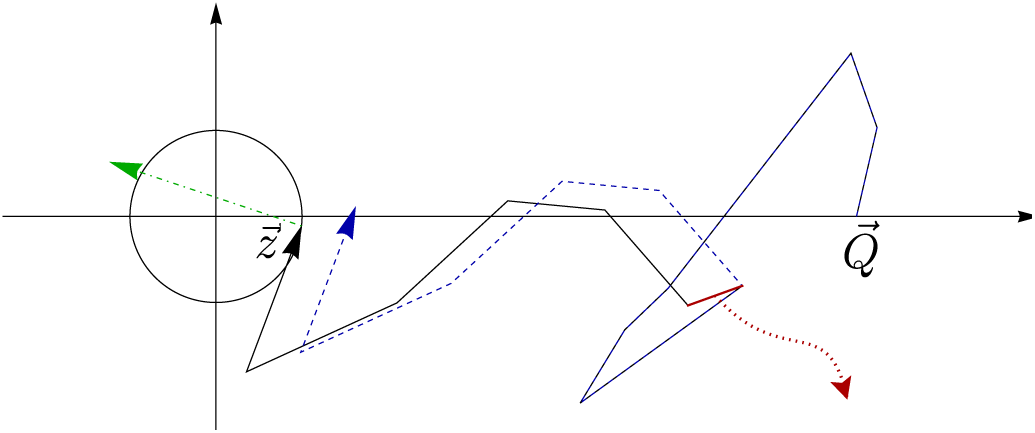
Illustration of the condition for a genotype to be a local fitness maximum. The black circle encloses phenotypes that have higher fitness than the focal phenotype 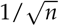. For *τ* to be a genotypic fitness maximum, both a phenotype with a further mutation (dash-dotted green arrow) and a phenotype without one of the mutations in *τ* (red segment and blue dotted arrows) should lie outside the circle.

### Summary 1

When the phenotypic dimension *n* is large and the Fisher parameter *x* is moderate, the probability of reciprocal sign epistasis decays as 1/*n*, while that of simple sign epistasis decays as 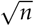. Although these probabilities decrease monotonically with *n* at fixed *x*, they have a non-monotonic behavior as a function of *x*: For small *x* they increase with *x* and for large *x* they decrease with *x* (see Figure 3). Under the pleiotropic scaling adopted in this work this implies that the probabilities are non-monotonic function of the wild type distance *Q* at fixed *n* and vice versa. In contrast, under the total effect model, where both the wild type distance *Q* and *x* scale as 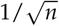, the probabilities decrease monotonically and exponentially with *n*.

## Genotypic complexity at fixed phenotypic dimension

In this section, we are interested in the number of local maxima in the genotypic fitness landscape. We focus on the expected number of maxima, which we denote by 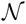, and analyze how this quantity behaves in the limit of large genotypic dimension, *L* → ∞, when the phenotypic dimension *n* is fixed. For the sake of clarity, the (unique) maximum of the phenotypic fitness landscape will be referred to as the phenotypic *optimum* throughout.probabilities decrease

### The number of local fitness maxima

Since fitness decreases monotonically with the distance to the phenotypic optimum, a genotype *τ* is a local fitness maximum if the corresponding phenotype defined by Equation 3 satisfies 
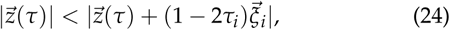
for all 1 ≤ *i* ≤ *L*. The phenotype vector appearing on the right hand side of this inequality arises from 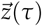 either by removing a mutation vector that is already part of the sum in Equation 3 (*τ_i_* = 1) or by adding a mutation vector that was not previously present (*τ_i_* = 0). The condition Equation 24 is obviously always fulfilled if 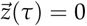, that is, if the phenotype is optimal, and we will see that in general the probability for this condition to be satisfied is larger, the more closely the phenotype approaches the origin. A graphical illustration of the condition Equation 24 is shown in Figure 5.

The ability of a phenotype 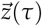 to approach the origin clearly depends on the number 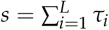 of mutant vectors it is composed of, and all phenotypes with the same number of mutations are statistically equivalent. The expected number of fit-with ness maxima can therefore be decomposed as

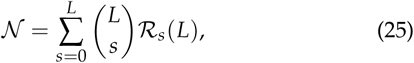
where 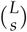 is the number of possible combinations of *s* out of *L* mutation vectors and 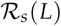 is the probability that a genotype with s mutations is a fitness maximum. The latter can be written as

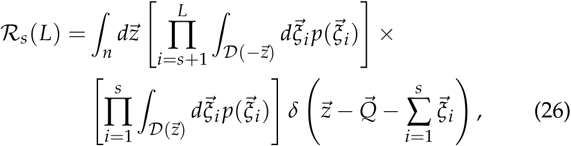
With 
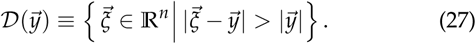

Here and below, ∫*_n_* stands for the integral over 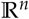.

Equation 26 can be understood as follows. First, the delta function 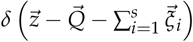 constrains 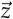 to be the phenotype of *τ* as defined in Equation 3. Next, the integration domains of the 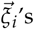 reflect the condition in Equation 24. Assuming without loss of generality that the *L* genetic loci are ordered such that *τ_i_* = 1 for *i* ≤ *s* and τ*_i_* = 0 for *i* > *s*, the maximum condition for *i* ≤ *s* requires 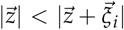, so the integration domain should be 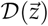, whereas for *i* > *s* the condition is 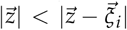, corresponding to the integration domain 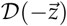. Using the integral representation of the delta function

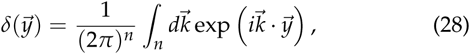
we can write 
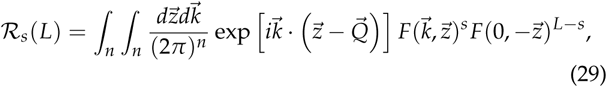
Where 
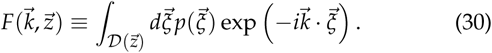

It was argued on qualitative grounds in **Model** that phenotypes that approach arbitrarily close to the origin are easily generated when the scaled wild type distance *q* is small, but they become rare for large *q*. As a consequence, it turns out that the main contribution to the integral over 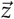 in Equation 29 comes from the region around the origin 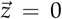 for small *q* but shifts to a distance *z* ∼ *L* along the 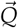-direction for large *q*. To account for this possibility, it is necessary to divide the integral domain into two parts 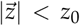 and 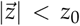, where *z*_0_ is an arbitrary non-zero number with *z*_0_/*L* → 0 as *L* → ∞. Thus, we write 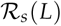 as

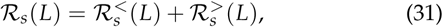

Where

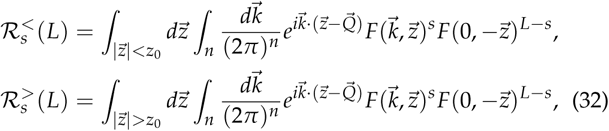
and correspondingly define 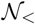 and 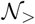 as

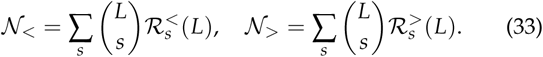

**Figure 6.**
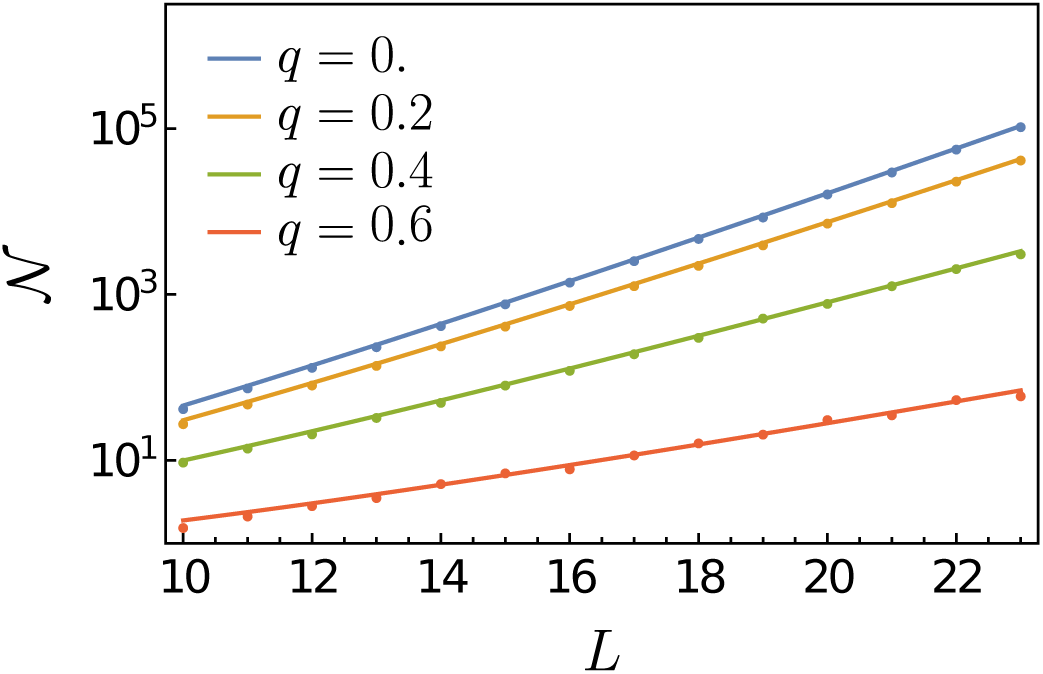
Plots of mean number of local maxima 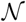 as a function of the genotypic dimension *L* for *q* **=** 0, 0.2, 0.4, and 0.6 with *n* = 1 on a semi-logarithmic scale. Data from numerical simulations are represented as dots, and the analytical prediction Equation 42 is shown as solid lines. Each dot represents the average over 10^5^ realizations of landscapes. In this parameter regime, **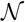** grows exponentially with *L* and the growth rate (i.e., the slopes of the lines) decreases with increasing *q*.

The total number of local maxima is then 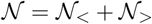.

***Regime I*.** We first consider 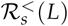. Expanding 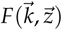 around the origin 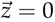, we show in Appendix C that

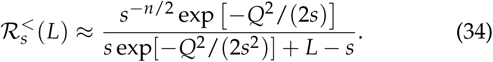

For an interpretation of Equation 34 it is helpful to refer to Figure 5. Note first that the probability that 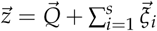 lies in the ball 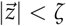 with radius ζ ≪ 1 is

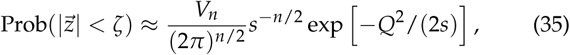
where *V_n_* (*ζ*) **∼** *ζ^n^* is the volume of the ball. We need to estimate how small ζ has to be for *τ* to be a local fitness maximum with an appreciable probability. Since the *s* random vectors contributing to 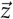 are statistically equivalent, it is plausible to assume that their average component parallel to 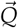 is 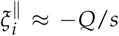 We further assume that the conditional probability density 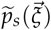 of these vectors, conditioned on their sum 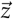 reaching the ball around the origin, can be approximated by a Gaussian, which consequently has the form 
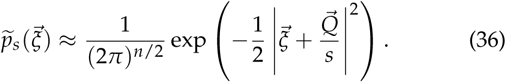

For 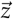 to be a phenotype vector of a local maximum, all these random vectors should lie in the region 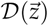 and the remaining (unconstrained) *L*—*s* vectors should lie in 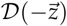. This event happens with probability

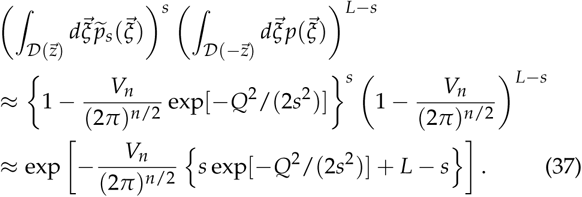

**Figure 7.**
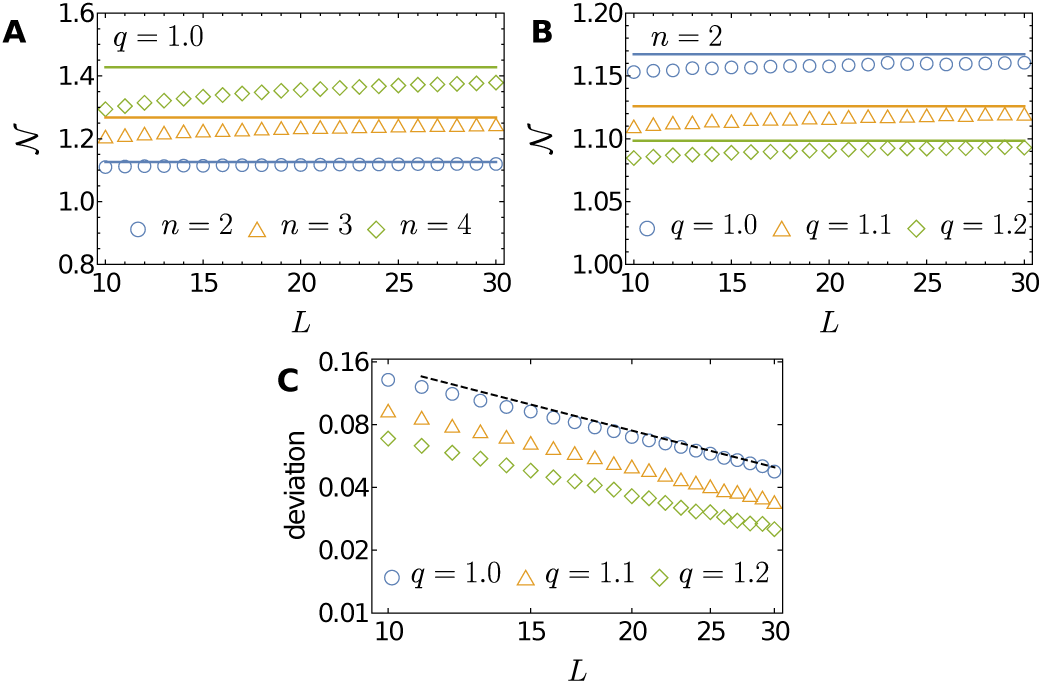
Comparison of simulation results (symbols) of the mean number of local maxima 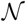 with analytic approximations (lines) for *q* > *qc*. Each symbol is the result of averaging over 2 x 10^6^ realizations. (**A**) 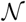 is shown to increase with *n* for fixed *q*. (**B**) 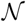 is shown to decrease with *q* for fixed *n*. (**C**) Deviation of the analytic expression from the simulation results, defined as 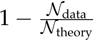, is depicted as a function of *L* on a double logarithmic scale. The phenotypic dimension for this panel is *n***=** 4, where the largest deviations are observed in panel **A**. The deviation decreases inversely with *L* as indicated by the black dashed line with slope -1.

Thus, we can estimate the typical value of ζ as the solution of

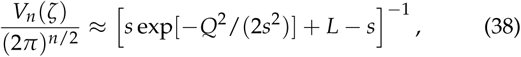
which, combined with Equation 35, indeed gives Equation 34.

To find the asymptotic behavior of 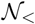 for large *L*, we use Stirling's formula in Equation 33 and approximate the summation over *s* by an integral over *ρ* ≡ *s*/*L*. This yields

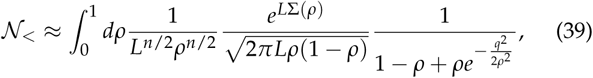
where the exponent Σ(*ρ*) is given by

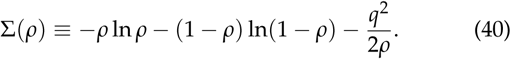

Under the condition *L* ≫ 1, the remaining integral with respect to *ρ* can be performed by expanding Σ(*ρ*) to second order around the saddle point *ρ*\* determined by the condition

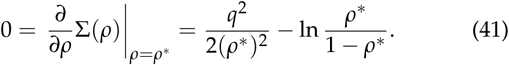

Performing the resultingGaussian integral with respect to *ρ* one finally obtains

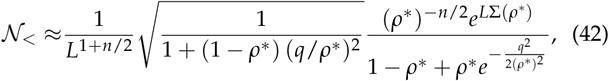
where *ρ* * = *ρ* * (*q*) is the solution of Equation 41, which is the (scaled) mean number of mutations in a local maximum. We will call *ρ***^*^** the mean genotypic distance. This solution is not available in closed form, but it can be shown that 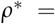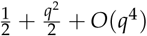 and Σ(*ρ*\*) = In2 – *q*^2^**+**O(*q*^4^) for small *q*. Figure 6 compares Equation 42 with the mean number of local maximum obtained by numerical simulations for various *q*'s with *n* = 1, to show an excellent agreement even for *L* = 10.

It is obvious that Σ(*ρ*) will eventually be negative as *q* increases for any value of *ρ*, and this must be true also for the maximum value Σ(*ρ*\*). Indeed, we found the threshold *q*_c_ ≈ 0.924 809, above which Σ(*ρ*\*) is negative. This signals a phase transition in the landscape properties. Inspection of Equation 40 shows that the transition is driven by a competition between the abundance of genotypes with a certain number of mutations and their likelihood to bring the phenotype close to the optimum. The first two terms in the expression for Σ(*ρ*) are the standard sequence entropy [see, for example, (Schimitt and Herzel 1997)] which is maximal at *ρ* = 1/2 (*s* = *L*/2), whereas the last term represents the statistical cost associated with *"*stretching*"* the phenotype towards to origin. With increasing *q*, the genotypes contributing to the formation of local maxima become increasingly atypical, in the sense that they contain more than the typical fraction *ρ* = 1/2 of mutations, and *ρ***^*^** increases. For *q* > *q*_c_ the cost can no longer be compensated by the entropy term and Σ(*ρ*^*^) becomes negative. In this regime 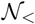 *decreases* exponentially with *L*, and therefore the total number of fitness maxima 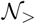, which by construction cannot be smaller than 1, must be dominated by the second contribution 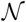.

***Regime II*.** We defer the detailed derivation of 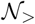 to Appendix C and here only report the final result obtained in the limit *L* → ∞, which is independent of *L* and reads

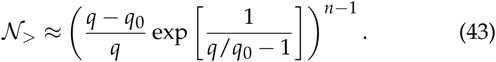

This expression is valid for 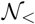, but it dominates the contribution 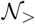 for large *L* only when *q* > *q*_c_. Figure 7 indeed shows that Equation 43 approximates the mean number of local maxima for *q* > *q*_c_, that is, 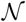 converges to 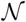 for large *L*. This figure also shows, as is clear by Equation 43, that 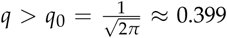 is a increasing (decreasing) function of *n* (*q*) for fixed value of *q* (*n*). The expected number of maxima is small in absolute terms in this regime, which can be attributed to the fact that the expression inside the parentheses in Equation 43 takes the value 1.214… at *q* = *q*_c_ and decreases rapidly towards unity for larger *q*.

To understand the appearance of *q*_0_, we refer to **Model**, where it was argued that 2*q*_0_*s* is the maximal distance towards the origin, which can be covered by a phenotype made up of *s* typical mutation vectors. Correspondingly, the analysis in Appendix C shows that the main contribution to 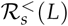 comes from phenotypes located at distance *z* = 2*s* (*q* – *q*_0_) from the origin, i.e., at distance 2s*q*_0_ from the wild type. The sum over *s* in Equation 33 is dominated by typical genotypes with *s* = *L*/2, and therefore the main contribution to 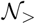 comes from phenotypes at distance *z* = (*q* - *q*_0_)*L* from the origin. The seeming divergence of 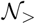 as 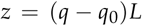 is an artifact of the approximation scheme, which assumes that the main contribution comes from the region where *z* ∼ O(*L*); clearly this assumption becomes invalid when 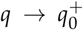. We note that for very large *q* and large *n* Equation 43 reduces to the expression 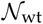 obtained in Equation 11 on the basis of Fisher’s formula for the fraction of beneficial mutations from the wild type phenotype.

**Figure 8.**
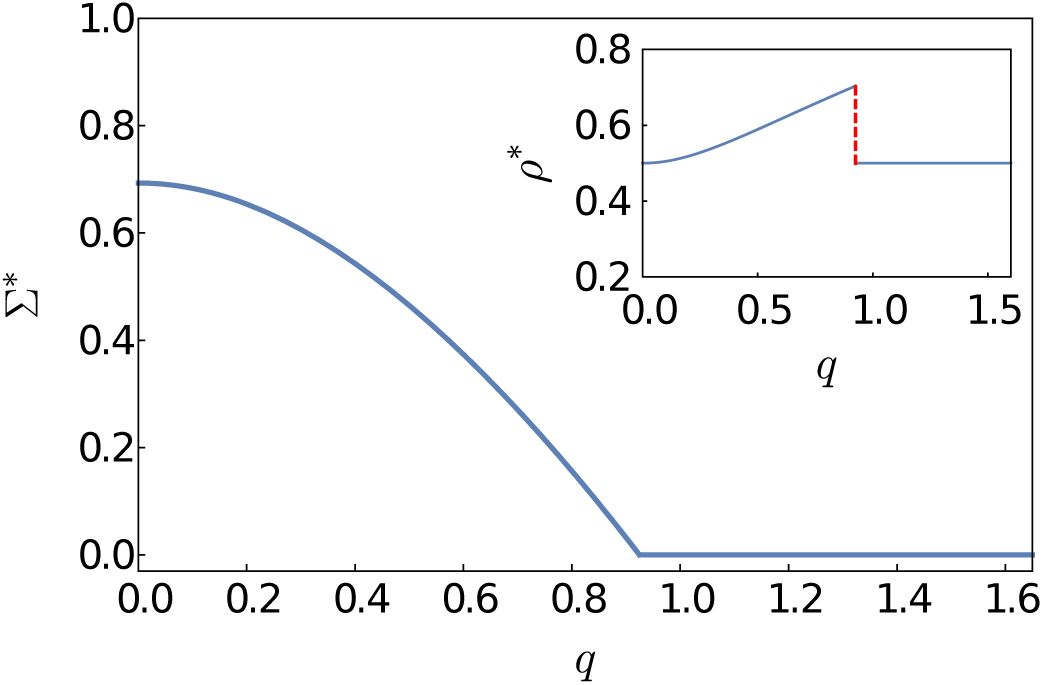
Plot of the genotypic complexity Σ* as a function of the scaled phenotypic wild type distance *q*. Here the phenotypic dimension *n* is kept finite while taking the genotypic dimension *L* to infinity. The complexity vanishes at the phase transition point *q* = *q*_c_ ≈ 0.924 809. Inset: Plot of the mean genotypic distance *ρ*^*^ of local maxima from the wild type as a function of *q*. Starting from 1/2, *ρ*\* increases with *q* for *q* < *q*_c_ and remains at 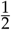 for *q* > *q_c_*.

### Phase transition

To sum up, the leading behavior of 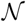 is

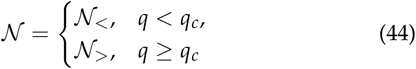
with 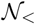 and 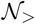 given by Equation 42 and Equation 43, respectively. Since 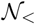 decreases to zero with *L* in a power-law fashion at *q* = *q_c_*, the dominant contribution at this value is 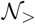. At *q* = *q_c_* the mean genotypic distance *ρ*\* jumps discontinuously from *ρ*\*(*q_c_*) ≈ 0.7035 to *ρ*\* = 1/2, and the mean phenotypic distance *z* * which is defined as the averaged magnitude of phenotype vectors for local maxima jumps from *z*\* ≈ 0 to *z*\* = (*q_c_* – *q_0_*)*L*. The genotypic complexity Σ* defined in Equation 13 is given by

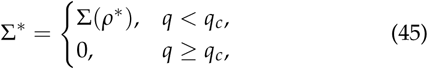
where *ρ*\* is the solution of Equation 41, and hence vanishes continuously at *q* = *q_c_*. These results are graphically represented in Figure 8. Recall that the value Σ* = ln2 attained at *q* = 0 is the largest possible, because the total number of genotypes is 2L = exp(*L* ln^2^). Remarkably, these leading order results are independent of the phenotypic dimension. A dependence on *n* emerges at the subleading order, and it affects the number of fitness maxima in qualitatively different ways in the two phases. For *q* < *q_c_* the pre-exponential factor in Equation 42 is a power law in *L* with exponent 1 + *n* /2 and hence decreases with increasing *n*, whereas the expression in Equation 43 describing the regime *q* > *q_c_* increases exponentially with *n*.

**Figure 9.**
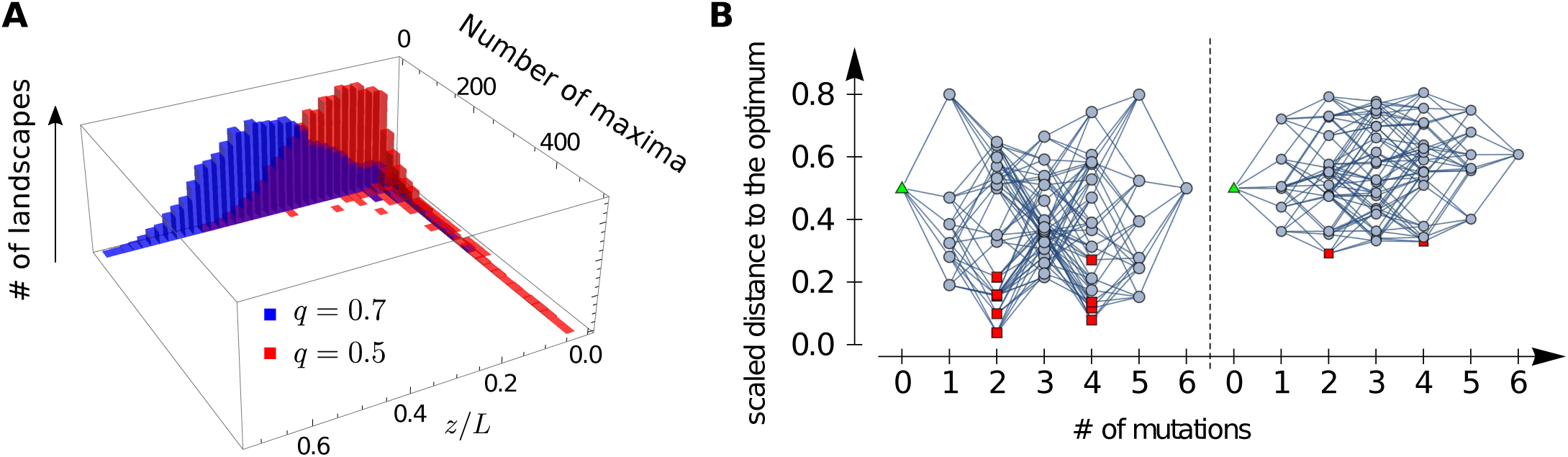
Coexistence of the two mechanisms I and II for *q*_0_ < *q* < *q*_c_. (**A**) Two dimensional histogram of the number of fitness maxima and the average phenotypic distance of the maxima to the optimum within a *single* realization. Here *L* = 15 and *n* = 2 are used and 10^4^ different landscapes are randomly generated for each value of *q*. Only a small number of realizations have a small average distance but these contribute an exceptionally large number of fitness peaks. (**B**) Two examples of genotype-phenotype maps selected from realizations with *q* = 0.5, *L* = 6, and *n* = 2. The wild type phenotype is marked by a green triangle and local fitness maxima by red squares. When the phenotypes of the local fitness maxima are close to (far away from) the origin, the number of maxima is large (small), which corresponds to mechanism I (II).

### Interpretation

The phase transition reflects a shift between two distinct mechanisms for generating genotypic complexity in FGM, which are analogous to the two origins of pairwise sign epistasis that were identified by Blanquart *et al*. (2014) and discussed above in ***Sign epistasis*.** In regime I (*q* < *q*_c_) the mutant phenotype closely approaches the origin and multiple fitness maxima are generated by overshooting the phenotypic optimum. By contrast, in regime II (*q* > *q*_c_) the phenotypic optimum cannot be reached and the genotypic complexity arises from the local curvature of the fitness isoclines. These two situations are exemplified by the two panels of Figure 1. For the sake of brevity, in the following discussion we will refer to the two mechanisms as mechanism I and mechanism II, respectively.

The approach to the origin in regime I is a largely onedimensional phenomenon governed by the components of the mutation vector along the direction of the wild type phenotype 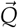, which explains why the leading order behavior of the genotypic complexity is independent of *n*. For *q* < *q*_c_, the *n*- dependence of the pre-exponential factor in Equation 42 arises from the increasing difficulty of the random walk formed by the mutational vectors to locate the origin in high dimensions. By contrast, mechanism II operating for *q* > *q*_c_ relies on the existence of the transverse dimensions, which is the reason why 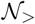 in Equation 43 is an increasing function of *n* with 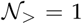 for *n* = 1.

When *q*_0_ < *q* < *q*_c_, both mechanisms seem to be present simultaneously. As our analysis is restricted to the average number of local maxima, at this point we cannot decide whether both mechanisms appear in a single realization of the fitness landscape, or if one of them dominates for a given realization. To answer this question, we generated 10^4^ fitness landscapes randomly for given parameter sets and identified all local maxima for each landscape. We then determined the number of local maxima and averaged the phenotypic distance of the local maxima to the optimum for each realization. This mean distance will be denoted by 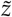 and is itself a random variable; it should not be confused with the mean phenotypic distance *z*\*, which is calculated by taking an average over all fitness peaks in all realizations, giving the same weight to each peak. The results are depicted as a two-dimensional histogram in Figure 9(A).

The figure shows that the marginal distribution of 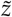 displays a pronounced peak around 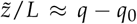, which corresponds to the behavior that is typical of mechanism I. For most realizations 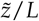 deviates significantly from zero, and only a small number of landscapes have local maxima near *z* = 0. However, these landscapes have many more maxima than typical landscapes and therefore dominantly contribute to the mean number of maxima 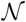. This shows that within a single realization the two mechanisms are not operative together and only a single mechanism exists. Since most realizations exhibit mechanism II, whereas the mean number of local maxima grows exponentially as expected for mechanism I, we conclude that mechanism I occurs rarely but once it does, it generates a huge number of local maxima, which compensates the low probability of occurrence. We may thus say that both mechanisms coexist for *q*_0_ < *q* < *q*_*c*_ and *q*_0_ can be regarded as the threshold of coexistence. Two fitness landscape realizations generated for the same value of *q* located in the coexistence region that exemplify the two mechanisms are shown in Figure 9(B).

### Summary 2

If the dimension *n* of phenotypic space is much smaller than the dimension *L* of genotypic space, there exists a threshold *q*_c_ of the scaled wild distance *q* to the phenotypic optimum below which the mean number 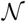 of local maxima in a genotypic fitness landscape increases exponentially with *L* and above which it saturates to a finite value. The genotypic complexity Σ*, which is defined as the exponential growth rate of 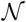 with *L*, is a decreasing function of *q* but does not depend on *n*. On the other hand, 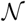 decreases with *n* for *q* < *q*_c_ yet increases with *n* for *q* > *q*_c_. Figure 8 depicts Σ^*^ and the mean genotypic distance *ρ*\* as functions of *q*. For *q*_0_ < *q* < *q*_c_, where 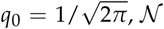 is dominated by a small fraction of landscape realizations that display an exceptionally large number of max-ima. If the pleiotropic scaling is assumed to follow the total effects model, we need to specify how the unscaled wild type distance *d* in Equation 2 depends on *L*. Assuming that *d* = *d_0_L* where *d_0_* is independent of *n* (Orr 2000), the scaled wild type distance 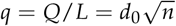 becomes an increasing function of *n*, and therefore the relation *q* < *q_c_* for regime I is never realized when *n* is sufficiently large.

**Figure 10.**
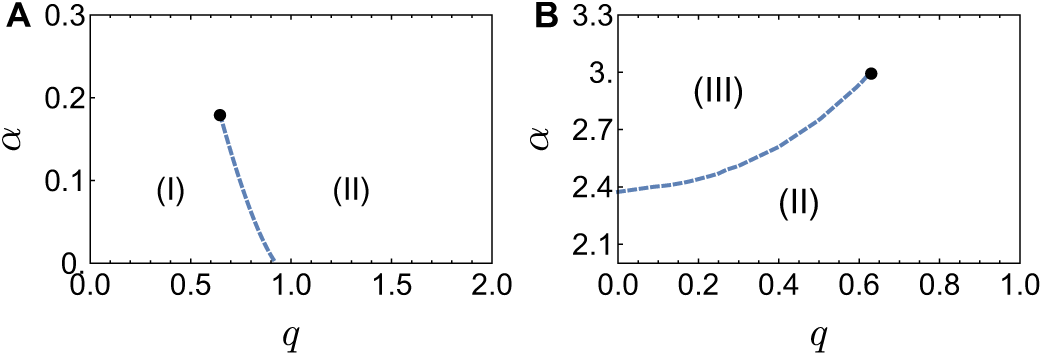
Phase diagrams in the parameter space (*q*,α). Here, *q* = *Q/L* is the scaled distance of the wildtype phenotype from the origin and *α* **=** *n***/** *L* is the ratio of phenotypic dimension to genotypic dimension. Dashed lines are phase boundaries at which the mean genotypic and phenotypic distances change discontinuously. (**A**) The phase boundary separating regimes I and II starts at (*q*,α) ≃ (0.925,0) and continues to exist until approximately *α* ≃ 0.18. (**B**) The phase boundary separating regimes II and III starts at (*q*, α) ≃ (0,2.38) and continues to exist until approximately *q* ≃ 0.62.

## Genotypic complexity in the joint limit

In the previous subsection, we have calculated the mean number of local fitness maxima 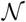 at a fixed phenotypic dimension *n*, assuming that the genotypic dimension *L* is much larger than *n*(*L* ≫ *n*). However, in applications of FGM one often expects that both *L* and *n* are large and possibly of comparable magnitude. In this case the results derived above can be unreliable for large *n*, as exemplified by the fact that the subleading correction to Equation 42 is of the order of O(*L^-1/n^*)(see Appendix C).

To obtain a reliable expression for 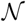 that is valid when both *n* and *L* are large, we now consider the joint limit *n, L* → ∞ at fixed ratio *α* = *n*/ *L*. This will allow us to find the leading behavior of the mean number of local maxima with a correction of order O(1/*L*). Furthermore, we will clarify the role of the phenotypic dimension in the two phases described in the previous subsection, and we will uncover a third phase that appears at large *α* (see Figure 10).

### The number of local fitness maxima

We relegate the detailed calculation to Appendix D and directly present our final expression for the mean number of local maxima,

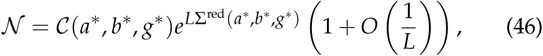
where the function Σ^red^ (*a, b, g*) in the exponent is given by

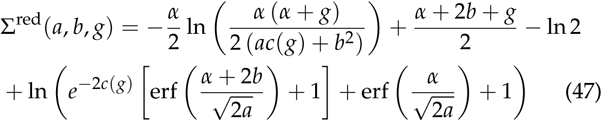
with 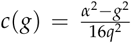 As before, the starred variables *a*, b\** and *g***^*^** denote the solution of the extremum condition

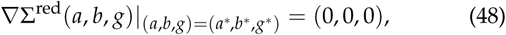
where ▽ is the gradient with respect to the three variables (*a, b, g*). When several solutions of Equation 48 exist, the one giving the largest value of Σ^red^ is chosen. The prefactor *C*(*a*, b***\*,** g*), which is independent of *L*, can be determined from Equation D17 presented in Appendix D. Even though the variables ( *a*, *b*, *g*) lack a direct intepretation in terms of the original setting of FGM, we show in Appendix E that *a***^*^** is related to the mean phenotypic distance *z\** by the equation 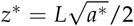.

**Figure 11.**
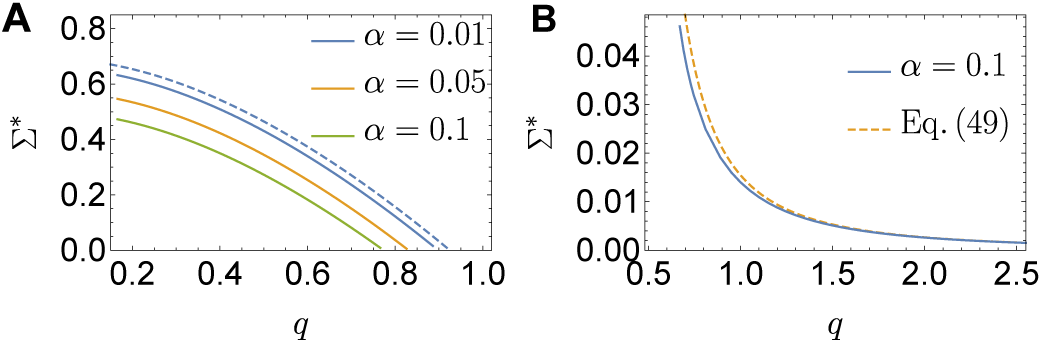
Convergence of the complexity to the fixed *n* case for small *α*. (**A**) The solid lines depict numerical solutions of Equation 48 for values of *α* belonging to regime I. The convergence to Equation 42 (dashed line) is clearly seen as *α*→ 0. (**B**) The blue solid line depicts the numerical solution of Equation 48 for *α***=** 0.1 belonging to regime II. Except for a slight devitation detectable when *q* is close to *q*_0_, Equation 49 (dashed line) remains a good approximation.

An immediate consequence of Equation 46 is that the number of local maxima increases exponentially in *L* for any value of *q* and *α* without algebraic corrections of the kind found in Equation 42. Obtaining closed form solutions of Equation 48, which ultimately determine the functional dependence of the complexity Σ^*^ on *α* and *q*, seems to be a formidable task. Instead, we resort to numerical methods by sweeping through the most interesting intervals, *q* ∈ (0, 2) and *α* ∈ (0, 3). Surprisingly, we find three independent branches of solutions that correspond to distinct phases. In order to acquire a qualitative understanding of these branches, it is instructive to first focus on the small *α* behavior, where one expects a smooth continuation to the results of Equation 42 and Equation 43 as *α*→ 0.

***Small*** *α* ***behavior*.** In contrast to the fixed *n* case where two separate analyses were carried out for the two regimes *q* < *q_c_* and *q* **>** *q_c_*, the present approach yields a single expression describing the genotypic complexity for arbitrary values of *q* and *α*. Consistently with the fixed *n* analysis, only two out of the three branches of solutions that were found in the numerical analysis exist for sufficiently small *α*, and they are separated by a phase transition as shown in the phase diagram in Figure 10(A). By extrapolating the behavior of Σ* towards *α* → 0 as shown in Figure 11, we are able to identify the correct counterparts for each of the two previously found regimes.

The extrapolation is straightforward in regime II, where the replacement *n* → *αL* in Equation 43 yields an exponential dependence of 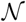 on *L* with the growth rate

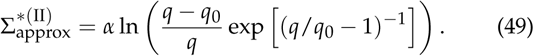

**Figure 12.**
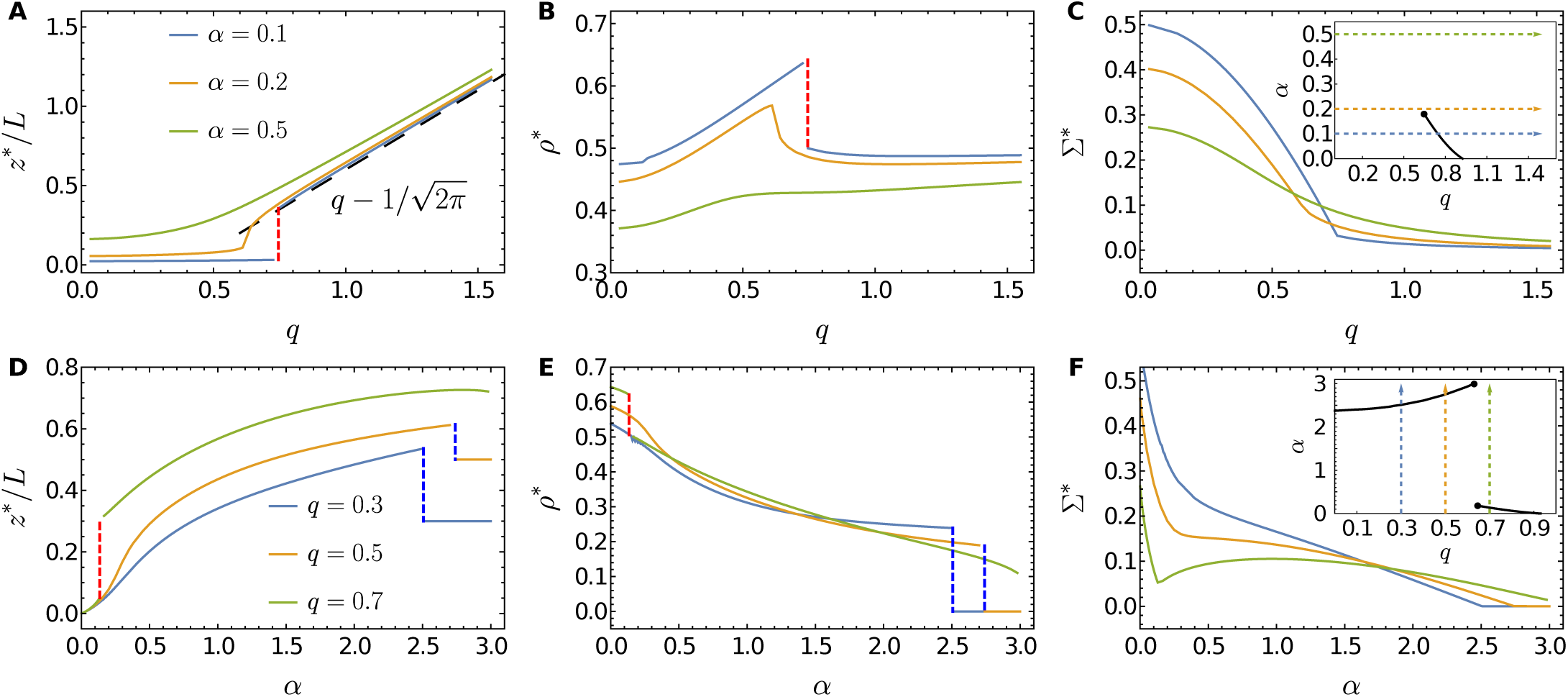
Plots of scaled mean phenotypic distance *z*\* /*L* (left column), mean genotypic distance *ρ*\* (middle column), and genotypic complexity Σ* (right column) against *q* for fixed *α* (top row) and against *α* for fixed *q* (bottom row). The curves in the top (bottom) panels are drawn along the arrows in the inset of panel **C** (**F**). Top row (**A,B,C**): When *α* is small, the landscape behaves similar to the fixed *n* case which effectively corresponds to *α* = 0. In this case *z*\* and *ρ*\* for large *q* are well approximated by *q* – *q*_0_ and 1/2, respectively. As *α* increases beyond the transition line, the first order transition visualized by the red dashed lines disappears and all quantities change smoothly with *q*. Bottom row (**D,E,F**): As *α* increases for small *q*, another phase transition with discontinuities in *z*^*^ and *ρ*^*^ (blue dashed lines) signals the appearance of regime III. The genotypic maxima in regime III are located very close to the wild type position, *z*\*/*L* ≃ *q* and *ρ*\* ≃ 0. This transition ceases to exist when *q* exceeds approximately 0.62. Note that the dependence of Σ* on *α* is non-monotonic for *q* = 0.7 (panel **F**).

This crude approximation turns out to be remarkably accurate even at *α* = 0.1, as illustrated in Figure 11(B). By contrast, in regime I the naive replacement of *n* by *αL* in Equation 42 yields an expression that vanishes faster than exponential in *L*, as exp[-(*α*/2)*L* ln*L*]. This reflects the fact that the mean phenotypic distance *z*^*^ moves away from the origin for any *α* > 0 and hence the complexity cannot be derived only by inspecting Equation 26 around *z* = 0 (see Figure 12 (A, D)). At the same time the mean genotypic distance *ρ*^*^ decreases with increasing *α* and eventually falls below the value *ρ*\* = 1/2 favored by the sequence entropy (Figure 12 (B, E)).

Both trends can be attributed to the increasing role of the perpendicular mutational displacements that make up the second term on the right hand side of Equation 5. Under the scaling of the joint limit, this term is of order *ρL*(*n* – 1) ≈ *ρ*α*L*^2^ and hence comparable to the first term originating from the parallel displacements. The perpendicular displacements always increase the phenotypic distance to the origin, and they are present even when *q* = 0. The additional cost to reduce the perpendicular contribution results in a smaller value of Σ^*^ compared to the case of fixed *n*. Moreover, whereas the parallel contribution is minimized (for *q* > *q*_0_) by making *ρ* as large as possible, the reduction of the perpendicular displacements requires small *ρ*.

In the fixed *n* analysis the number of fitness maxima was found to decrease (increase) with *n* in regime I (II) and this tendency is recovered from the joint limit case when *α* is not too large (Figure 12 (C)). Because of these opposing trends of Σ* in the two regimes, the location of the phase transition separating them is expected to decrease with increasing *α*, as can be seen in Figure 10(A). If one ignores the contribution from the perpendicular displacements, the phenotypic position of the fitness maxima is expected to jump from *z*^*^ = 0 to *z*^*^ = *q*_*c*_ – *q*_0_ at the transition, and thus the jump size should decrease as *q*_c_ decreases. This observation suggests that the two branches should merge into one when *q*_*c*_ reaches *q*_0_. With the additional contribution of perpendicular dimensions, we numerically found that this critical end point at which the phases I and II merge occurs even earlier, at *α* ≃ 0.18 and *q* ≃ 0.62 > *q*_0_ (Figure 10). For *α*larger than 0.18, *ρ*\* does not show any discontinuity for any *q* as long as the parameters are in the regime II.

***Large*** *α* ***behavior and regime III*.** In order to develop some intuition about the FGM fitness landscape in the regime where *α* = *n*/*L* ≫ 1, we revisit the results obtained in ***Sign epistasis*,**where pairs of mutations were considered. Two conclusions can be drawn about the typical shape of these small genotypic landscapes (of size *L* = 2) in the limit *n* → ∞. First, the probability that the wild type is a genotypic maximum tends to unity according to Equation 10. Second, the joint distribution given in Equation 15 enforces additivity of mutational effects for large *n*, and correspondingly the probability for sign epistasis vanishes. Thus for large *n* the two-dimensional genotypic landscape becomes smooth with a single maximum located at the wild type. Assuming that this picture holds more generally whenever the limit *n* → ∞ is taken at finite *L*, we expect the following asymptotic behaviors of the quantifiers of genotypic complexity for large *α*: (i) 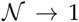, Σ* → 0 (unique genotypic optimum); (ii) *z*\*/*L* → *q, ρ*\* → 0 (location of the maximum at the wild type phenotype and genotype).

This expectation is largely borne out by the numerical results shown in the bottom panels of Figure 12. However, depending on the value of *q*, the approach to the limit of a smooth landscape can be either continuous (for large *q*) or display characteristic jumps indicated by the blue dashed lines in Figure 12 (D) and (E). These jumps as well as the discontinuity in the slope of Σ^*^ as a function of *α* in Figure 12 (F) are hallmarks of the phase transition to the new regime III, which is represented by the dashed line in Figure 10 (B).

Fortunately, the solution of Equation 48 describing the new phase can be obtained analytically from Equation 47 or Equation F3 as a series expansion. The derivation presented in Appendix G yields

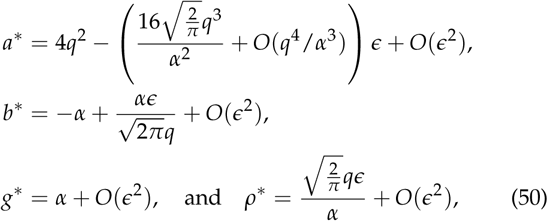
where the expansion parameter 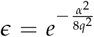 decays rapidly with increasing *α****/****q*. The corresponding genotypic complexity can also be evaluated in a series expansion,

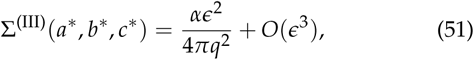
which shows that Σ^*^ is positive but vanishingly small in this regime. We note that using Equation E5, the expression for *a*^*^ in Equation 50 amounts to

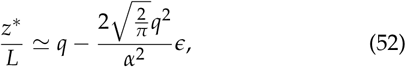
implying that the small number of local maxima that exist in this phase are located very close to the wild type phenotype.

To first order in ∈, the results for *ρ*\* and *z*\* in Equation 50 and Equation 52 can be easily derived from the idea that mutational effects become approximately additive for large *α*, thus providing further support for this assumption. If mutational effects are strictly additive, the probability for a genotype contaning *s*>mutations to be a local fitness maximum is given by

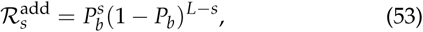
where *P_b_* is the probability for a mutation to be beneficial. Equation 53 expresses the condition that reverting any one of the *s* mutations contained in the genotype as well as adding one of the unused *L* – *s* mutations should lower the fitness. Using Fisher’s Equation 1, the probability for a beneficial mutation is 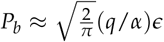 for large *α*. Thus to linear order in ∈ or *P_b_*, the expected number of mutations contributing to such a genotype is 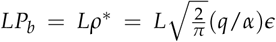, which is consistent with Equation 50.

The phenotypic location of a local maximum deviates from 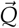 in those rare instances where one of the mutations from the wild type is beneficial, which happens with probability *P_b_*. To estimate the corresponding shift in *z*\*, we refer to the results of subsection **Sign epistasis**, where it was shown that the squared phenotypic displacement *R*_1_ defined in Equation 14 has a Gaussian distribution with mean 1 and variance 1/*x*^2^ = 4*q*^2^ / α^2^ for large *n*. Using this, it is straightforward to show that the expected value of *R*_1_ conditioned on the mutation to be beneficial (*R*_1_ < 0) is 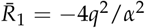 to leading order. Multiplying this by the expected number of mutations *LP_b_* we obtain the relation

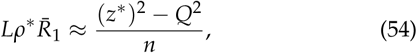
which yields the same leading behavior for *z*^*^ / *L* as in Equation 52.

**Figure 13.**
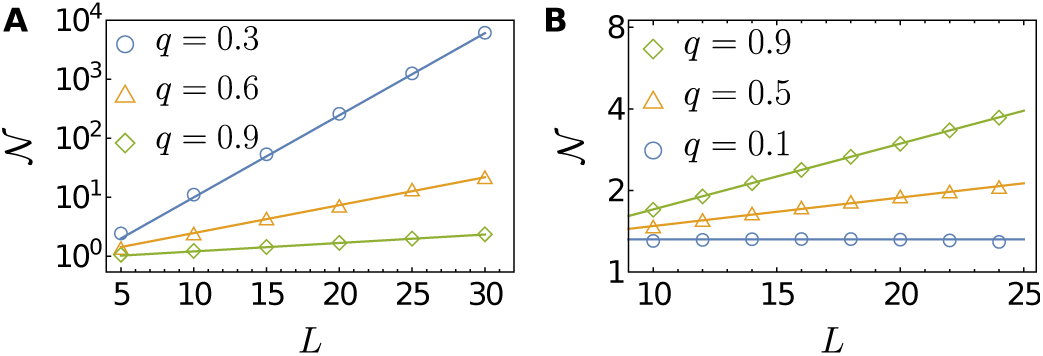
Semi-logarithmic plots of the mean number of local maxima 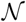 vs. the genotypic dimension *L* for (**A**) *α* = 0.2 and (**B**) *α* = 2.5 and for various values of *q*. Each symbol represents the average over 10^5^ randomly generated landscapes, and lines depict the analytic approximation of Equation D17. The approximation is good even for moderate *L*.

As previously observed for the transition between regimes I and II, the phase boundary separating regimes II and III terminates at a point where the two solutions defining the regimes merge (Figure 10 (B)). Beyond this point the jumps in *z*^*^ and *ρ\** seen in Figure 12 (D) and (E) disappear and all quantities approach smoothly to their asymptotic values. A surprising feature of the large *α* behavior that persists also for larger *q* is that the complexity becomes an *increasing* function of *q* when *α* > 1.7 (Figure 12 (F)). In Figure 13 we verify this behavior using direct simulations of FGM. These simulations also show that the predictions based on Equation 47 are remarkably accurate already for moderate values of *L* and *n*.

***Summary 3*.** When the dimension *n* of the phenotypic trait space and the dimension *L* of the genotypic space are large and comparable, the genotypic complexity Σ^*^ is always nonzero and depends on the ratios *α* = *n*/ *L* and *q* = *Q***/** *L*. There are three regimes where the behavior of the genotypic complexity and the mean genotypic distance *ρ*^*^ (the average number of mutations in a local maximum divided by *L*) are qualitatively different. In regime I which is roughly characterized by small *q* and small *α*, there are many local maxima in the region located far away from the wild type but close to the phenotypic optimum, and the fitness landscape is quite rugged. In regime II which is roughly characterized by large *q* and small *α*, there is an appreciable number of local maxima, though smaller than in regime I, and typically half of the *L* mutations contribute to the corresponding genotypes. In regime III which is roughly characterized by large *α*, the genetic complexity is very small, though nonzero. Also *ρ*^*^ is close to zero, which means that the wild type has a high probability to be the global fitness maximum. An overview of the three regimes is found in Table 2.

## Discussion

Fisher’s geometric model (FGM) provides a simple yet generic scenario for the emergence of complex epistatic interactions from a nonlinear mapping of an additive, multidimensional phenotype onto fitness. Its role in the theory of adaptation may be aptly described as that of a *"*proof of concept model*"* (Servedio *et al*. 2014), and as such it is widely used in fundamental theoretical studies (Blanquart *et al*. 2014; Chevin *et al*. 2010; Fraïsse *et al*. 2016; Gros *et al*. 2009; Martin 2014; Moura de Sousa *et al*. 2016) as well as for the parametrization and interpretation of empirical data (Bank *et al*. 2014; Blanquart and Bataillon 2016; Martin *et al*. 2007; Perfeito *et al*. 2014; Schoustra *et al*. 2016; Velenich and Gore 2013; Weinreich and Knies 2013). Rather than tracing the mutational effects and their interactions to the underlying molecular basis, the model aims at identifying robust features of the adaptive process that can be expected to be shared by large classes of organisms.

**Table 2.** Characteristics of the three regimes in the joint limit

| Regime | Condition | $\Sigma^*$ | $\rho^*$ | landscape |
| --- | --- | --- | --- | --- |
| I | $q \ll 1, \alpha \ll 1$ | $> 0$ | $> 1/2$ | rugged |
| II | $q \gg 1, \alpha \ll 1$ | $\simeq 0$ | $\approx 1/2$ | intermediate |
| III | $\alpha \gg 1$ | $\simeq 0$ | $\simeq 0$ | almost smooth |

To give an example of such a feature that is of central importance in the present context, it was pointed out by Blanquart *et al*. (2014) that pairwise sign epistasis is generated in FGM through two distinct mechanisms. In one case the mutational displacements overshoot the phenotypic optimum, whereas in the other case the displacements are directed approximately perpendicular to the direction of the optimum and sign epistasis arises because the fitness isoclines are curved. The first mechanism is obviously operative also in a one-dimensional phenotype space, but in the second case [termed antagonistic pleiotropy by Blanquart *et al*. (2014)] at least two phenotypic dimensions are required. Interestingly, both mechanisms have been invoked in empirical studies where a nonlinear phenotype-fitness map was used to model epistatic interactions between multiple mutations. In one study, Rokyta *et al*. (2011) explained the pairwise epistatic interactions between 9 beneficial mutations in the ssDNA bacteriophage ID11 by assuming that fitness is a single-peaked nonlinear function of a one-dimensional additive phenotype. In the second study the genotypic fitness landscapes based on all combinations of two groups of four antibiotic resistance mutations in the enzyme *β*-lactamase were parametrized by a nonlinear function mapping a two-dimensional phenotype to resistance (Schenk *et al*. 2013). The fitted function was in fact monotonic and did not possess a phenotypic optimum, which makes it clear that the epistatic interactions arose solely from antagonistic pleiotropy in this case.

In this work we have shown that the two mechanisms described by Blanquart *et al*. (2014) lead to distinct regimes or phases in the parameter space of FGM, where the genotypic fitness landscapes display qualitatively different properties (Figure 10(A)). When the phenotypic dimension *n* is much smaller than the genotypic dimension *L*, the two regimes are separated by a sharp phase transition where the average number and location of genotypic fitness maxima changes abruptly as the distance *q* of the wild type phenotype from the optimum is varied. In regime I (*q* < *q*_c_) the phenotypic optimum is reachable at least by some combinations of mutational displacements. Overshooting of the optimum is therefore possible and sign epistasis is strong, leading to rugged genotypic landscapes with a large number of local fitness maxima that grows exponentially with *L*. By contrast, in regime II (*q* > *q*_c_) only antagonistic pleiotropy is operative and the number of fitness maxima is much smaller. More precisely, for finite *n* the number tends to a finite limit for *L* → ∞, but the limiting value is an exponentially growing function of *n*.

**Figure 14.**
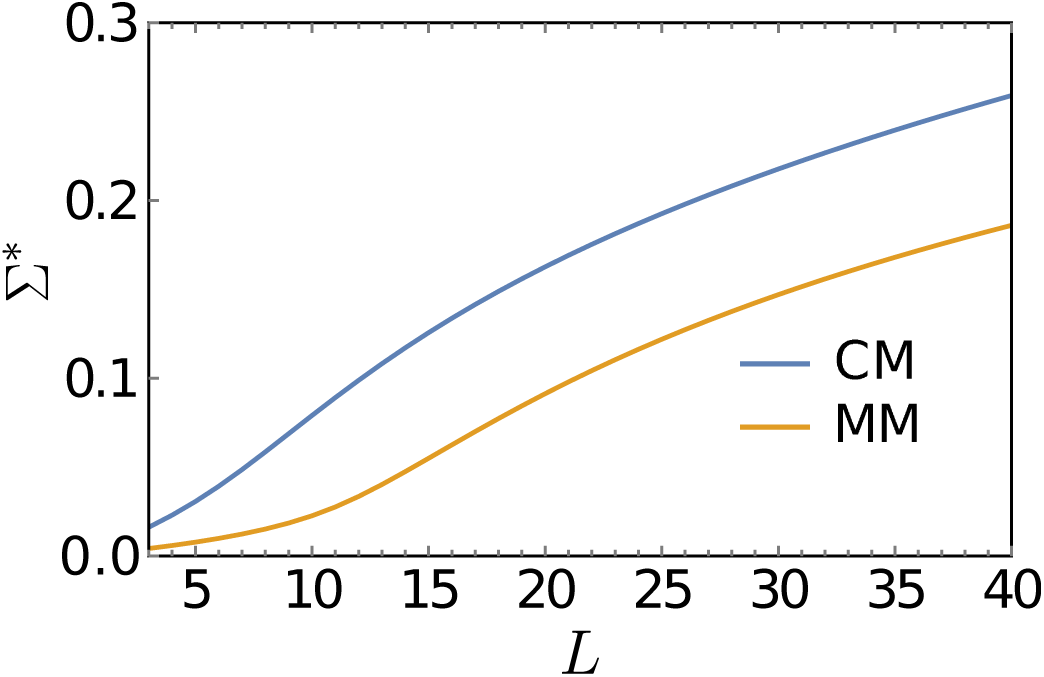
The logarithm of the number of local fitness maxima divided by the number of loci *L* is shown as a function of *L* for FGM with the parameter values *n* = 19.3, *Q* = 6.89 and *n* = 34.8, *Q* = 9.81 obtained by Schoustra *et al*. (2016) for the fungus *A. nidulans* growing in complete (CM) and minimal medium (MM), respectively. For the evaluation of 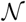 Equation 47 was used.

An important consequence of our results is that the dependence of the fitness landscape ruggedness on the phenotypic dimension *n* is remarkably complicated. For *n* ≪ *L*, landscapes become less rugged with increasing *n* in regime I (*q* < *q*_*c*_) but display increasing ruggedness in regime II (*q* > *q*_*c*_). When *n* ≫ *L* the ruggedness decreases with *n* for all *q* and the landscapes become approximately additive (regime III). In particular, the probability of sign epistasis vanishes algebraically with *n* in this regime. Thus *n* cannot in general be regarded as a measure of "phenotypic complexity", as a larger value of *n* does not imply that the corresponding fitness landscape is more complex.

This observation is relevant for the interpretation of experiments where the parameters of FGM are estimated from data. In recent work, FGM was used to analyze data on pairwise epistasis between beneficial mutations in the filamentous fungus *Aspergillus nidulans* growing in two different media (Schoustra *et al*. 2016). The estimates obtained for the phenotypic dimension and the distance of the wildtype phenotype from the optimum were *n* = 19.3, *Q* = 6.89 in complete medium and *n* = 34.8, *Q* = 9.81 in minimal medium, which, surprisingly, may seem to suggest a higher "phenotypic complexity" in the minimal medium. Using the results derived in this paper we can translate the estimated parameter values into the average number of maxima that a genotypic fitness landscape of a given dimension *L* would have. As can be seen in Figure 14, with respect to this measure the fitness landscape of the fungus growing in complete medium is actually more rugged. This is consistent with experiments using *Escherichia coli*, which found a greater heterogeneity of fitness trajectories in complete medium (Rozen *et al*. 2008), and indicates that the complete medium allows for a greater diversity of paths to adaptation than the minimal medium.

We hope that the results presented here will promote the use of FGM as part of the toolbox of probabilistic models that are currently available for the analysis of empirical fitness landscapes (Bank *et al*. 2016; Blanquart and Bataillon 2016; de Visser and Krug 2014; Hayashi *et al*. 2006; Neidhart et al.2014; Szendro *et al*. 2013). Compared to purely genotype-based models such as the NK and Rough-Mount-Fuji (RMF) models, FGM is arguably more realistic in that it introduces an explicit phenotypic layer mediating between genotypes and fitness (Martin 2014). Somewhat similar to the RMF model, the fitness landscapes of FGM are anisotropic and display a systematic change of properties as a function of the distance to the optimal phenotype (FGM) or the reference sequence (RMF), respectively (Neidhart *et al*. 2014). The idea that fitness landscape ruggedness increases systematically and possibly abruptly when approaching the optimum has been proposed previously in the context of *in vitro* evolution of proteins (Hayashi *et al*. 2006). If this is indeed a generic pattern, it may have broader implications. For example, de Visser *et al*. (2009) showed that the evolutionary benefits of recombination are severely limited by the presence of multiple peaks. If such peaks are rare far away from the optimum, the benefits of recombination would be most pronounced for particularly maladapted populations.

A recent investigation of 26 published empirical fitness landscapes using Approximate Bayesian Computation concluded that FGM could account for the full structure of the landscapes only in a minority of cases (Blanquart and Bataillon 2016). One of the features of the empirical landscapes that prevented a close fit to FGM was the occurrence of sign epistasis far away from the phenotypic optimum. Our analysis confirms that this is an unlikely event in FGM, and precisely quantifies the corresponding probability through Equation 16 and Equation 17. Blanquart and Bataillon (2016) also found that the phenotypic dimension is particularly difficult to infer from realizations of genotypic fitness landscapes, which matches our observation that the structure of the landscape depends only weakly on *n* when *n* ≪ *L*. We expect that our results will help to further clarify which features of an empirical fitness landscape make it more or less amenable to a phenotypic description in terms of FGM or some generalization thereof.

We conclude by mentioning some open questions that should be addressed in future theoretical work on FGM. First, a significant limitation of our results lies in their restriction to the *average* number of local fitness maxima. The number of maxima induced by a given realization of mutational displacements is a random variable, and unless the distribution of this variable is well concentrated, the average value may not reflect the typical behavior. The large fluctuations between different realizations of fitness landscapes generated by FGM were noticed already by Blanquart *et al*. (2014) on the basis of small-scale simulations, and they clearly contribute to the difficulty of inferring the parameters of FGM from individual realizations that was reported by Blanquart and Bataillon (2016). In the light of our analysis this pronounced heterogeneity can be attributed to the existence of multiple phases in the model, and it is exemplified by the simulation results in Figure 9. To quantitatively characterize the fluctuations between different realizations a better understanding of the distribution of the number of fitness maxima and its higher moments is required.

Second, the consequences of relaxing some of the assumptions underlying the formulation of FGM used in this work should be explored. The level of pleiotropy can be reduced by restricting the effects of mutational displacements to a subset of traits (Chevin *et al*. 2010; Moura de Sousa *et al*. 2016), and it would be interesting to see how this affects the ruggedness of the fitness landscape. However, the most critical and empirically poorly motivated assumption of FGM is clearly the absence of epistatic interactions on the level of phenotypes. It would therefore be important to understand how robust the results presented here are with respect to some level of phenotypic epistasis, which should ideally arise from a realistic model of phenotypic networks (Martin 2014).

Third, a natural extension of the present study is to consider multi-allelic genetic sequences. An immediate generalization keeping the additivity of mutational effects on the level phenotypes is to consider the following genotype-phenotype map

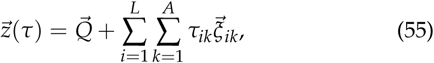
where *A* is the size of the alphabet from which the sequence elements are drawn (e.g., *A***=** 4 for DNA or RNA and *A***=** 20 for proteins), τ*_ik_* = 1 (0) if the allele at site *i* is (is not) *k*, and the 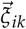 are uncorrelated random vectors. Clearly, our results for pairwise epistasis remain the same for this generalized model because they only concern mutations at different sites. However, the condition for a local fitness maximum now involves mutations to different alleles at the same site, which may lead to a non-trivial dependence on *A*. On the basis of a recent study of evolutionary accessibility in multi-allelic sequence spaces one may expect the fitness landscapes to become less rugged with increasing *A* (Zagorski *et al*. 2016), but this conjecture would have to be corroborated by a detailed analysis.

Finally, whereas the present work focused on the structure of the fitness landscapes induced by FGM, it is of obvious importance to understand how the adaptative process actually proceeds on such a landscape (Orr 2005). A simple framework that allows to address this question is provided by adaptive walks following Gillespie's strong selection/weak mutation dynamics (Gillespie 1983, 1984; Orr 2002). In a pioneering study, Orr (1998) considered adaptive walks in FGM assuming that the number *L* of possible mutations is unlimited. In this setting, any population not located precisely at the phenotypic optimum has a nonzero probability of generating another beneficial mutation and the adaptive walker never stops; see Park and Krug (2008) for a related analysis of adaptation in the house-of-cards landscape. For finite but large *L*, an interesting question concerns the number of steps until the population finds a local fitness maximum when the adaptive dynamics is random (Kauffman and Levin 1987; Park *et al*. 2015; Park and Krug 2016) or greedy (Orr 2003; Park *et al*. 2016). This problem is currently under investigation.

## Footnotes

1 When counting the number of local genotypic maxima, we checked all genotypes and counted the number exactly for a randomly realized landscape, then took an average.

## Acknowledgments

We thank Anton Bovier, David Dean, Guillaume Martin, Olivier Tenaillon and an anonymous reviewer for helpful remarks. This work was supported by DFG within SFB 680 *Molecular Basis of Evolutionary Innovations* and SPP 1590 *Probabilistic Structures in Evolution*. S-CP acknowledges the support by the Basic Science Research Program through the National Research Foundation of Korea (NRF) funded by the Ministry of Science, ICT and Future Planning (Grant No. 2014R1A1A2058694). S-CP would also like to thank Korea Institute for Advanced Study (KIAS) for its support and hospitality during his stay there on sabbatical leave (2016-2017).

## Appendix A: Derivation of the joint probability density 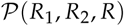

For the purpose of this calculation it will turn out to be convenient to locate the wild type phenotype on the diagonal of the trait space, i.e. to set 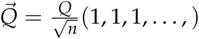. The probability density 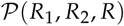 can then be formally defined as

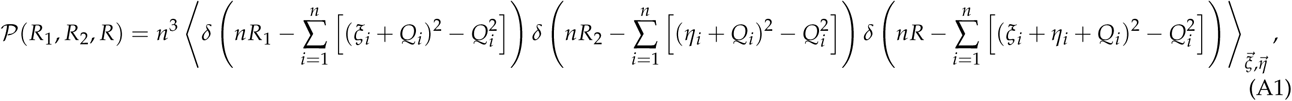
where 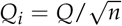 and 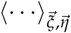 stands for the average over the distribution of 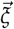 and 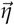 Using the integral representation of the delta function, we can write

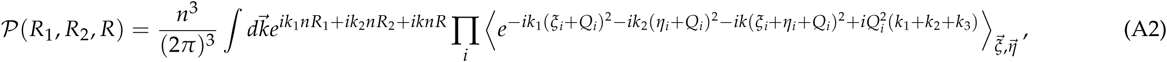
where 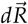 and 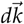 stand for *dR*_1_ *dR*_2_ *dR* and *dk*_1_ *dk*_2_ *dk*, respectively and we factorized the average by taking into account that the ξ*_i_****'***sand η*_i'_*s are all independent and identically distributed. The average in Equation A2 is readily calculated as

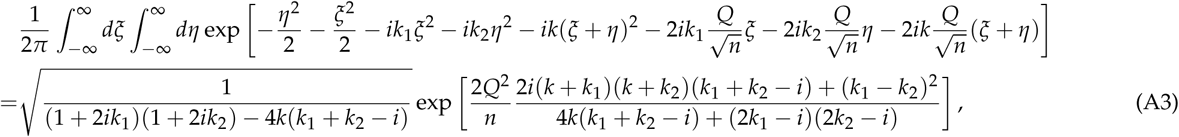
which gives

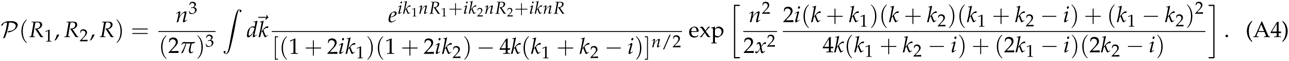

In the limit *n* → ∞, the integral is dominated by contributions from the vicinity of the extremum of the exponent, which can be algebraically determined to be *k* = *k*_1_ = *k*_2_ = 0. By expanding the argument of the exponential function up to the second order around this point and performing the Gaussian integral, we obtain

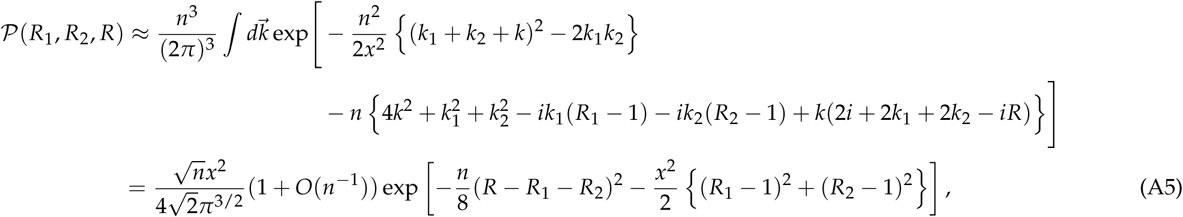
which is Equation 15.

## Appendix B: Probability of sign epistasis

In this appendix, we present the mathematical details of the derivation of the probabilities *P_r_* and *P_s_* of observing reciprocal sign epistasis (RSE) and simple sign epistasis (SSE), respectively. As in the main text, let us assume *R*_1_ < *R*_2_. In calculating the probabilities, the integral over *R* takes one of three forms

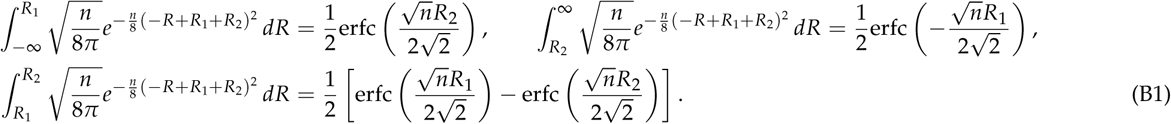

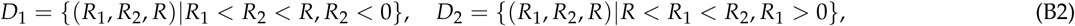
as illustrated in Figure 2. The probability of being in *D*_1_ is

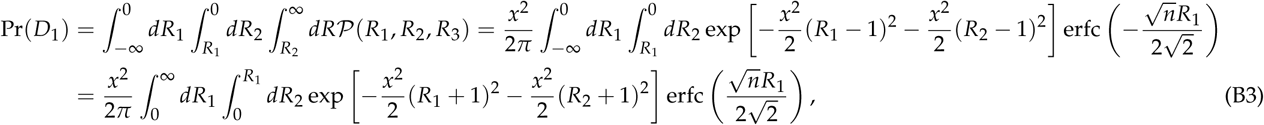
where we have changed variables *R_i_* → − *R_i_*. Since 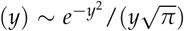 for *y*≫1, the above integral is dominated by the region *R*_1_ ≪ 1 for large *n*. Thus, it is sufficient to approximate 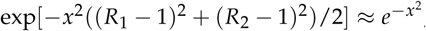, which yields

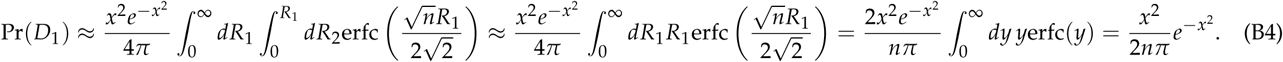

The probability of being in *D*2 has the same leading behavior,

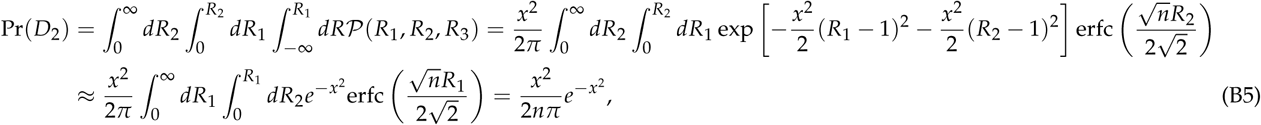
where we have exchanged the variables *R*_1_ ↔ *R*_2_. Due to the symmetrical roles of *R*_1_ and *R*_2_, the total probability of RSE is

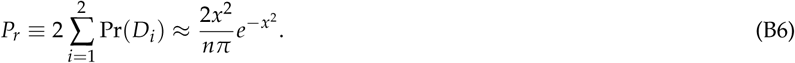

We can use a similar approximation scheme to calculate the probability of SSE. There are four domains contributing to SSE (see Figure 2),

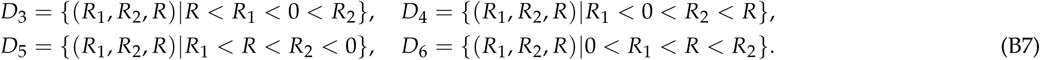

As we will see, all integrals can be represented by the functions

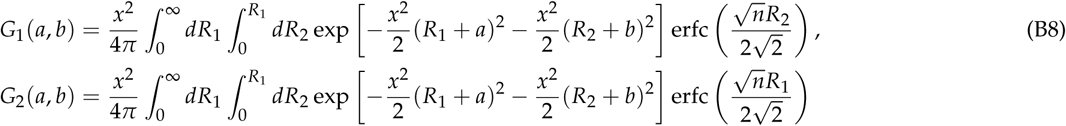

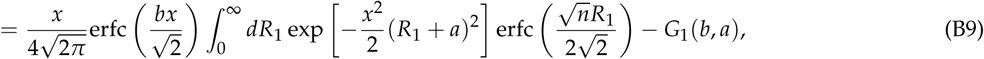
where *a, b* = ±1 and we have used that

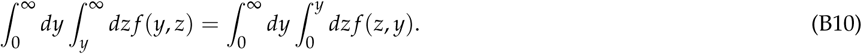

To be specific, we write the probabilities of being in each domain as

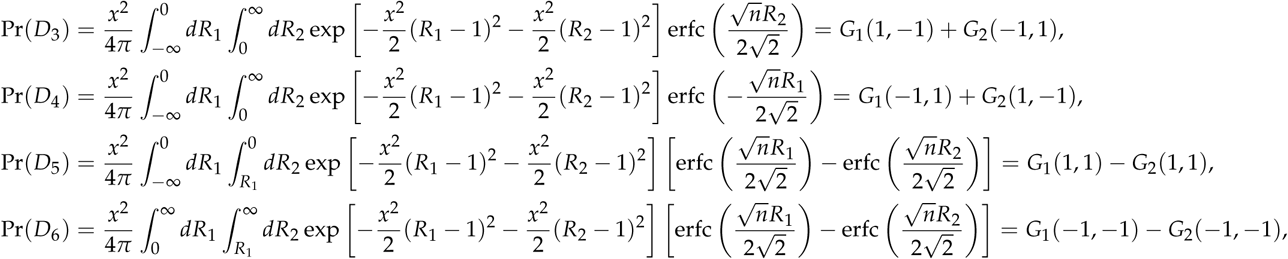
where we have changed negative integral domains into positive domains and made use of Equation B10. Using the approximation scheme explained above, we get

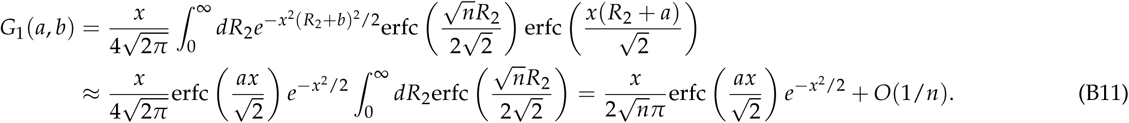

Since

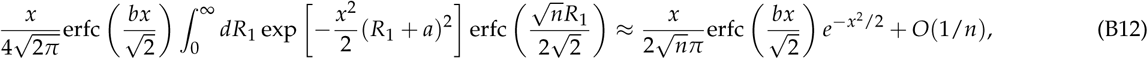
we conclude that *G*_2_(*a, b*) = O(1/*n*). Using erfc(*y*) + erfc(—*y*) = 2, we finally obtain

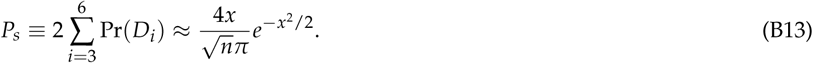

## Appendix C: Large *L* behavior of 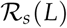 for fixed *n*

In this appendix, we calculate the asymptotic behavior of the probabilit 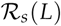 for a genotype with s mutations to be a local fitness maximum in the limit where *L* is large and the phenotype dimensions *n* is fixed. As explained in the main text, this probability has two contributions which arise from expanding the function 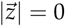 defined in Equation 30 near 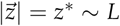 and 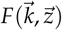, respectively (see Equation 31).

First, we consider the contribution from the region 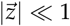. In this case, we can approximate 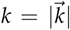 as

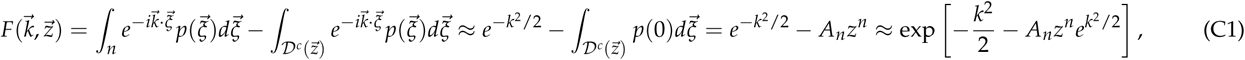
where 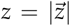, 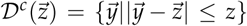,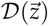 which is the complement of 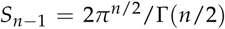, *A*_n_ = *p*(0)*S_n_*_-1_/*n* with 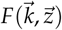 being the surface area of the unit sphere in (*n* - 1) dimensions, and *p*(0) = (2π)^—*n/2*^. Note that the error of the above approximation is O(*z*^*n+1*^). Thus, setting *ρ* ≡ *s*/*L* we can approximate

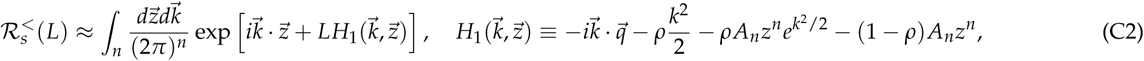
where 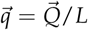. Since *L* is large, we can employ the saddle point approximation. One can easily see that the saddle point solving the equations 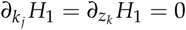 is at 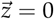 and *k_j_* = – *iq_j_*/ρ. Around the saddle point, we expand

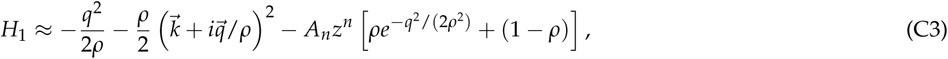
which gives

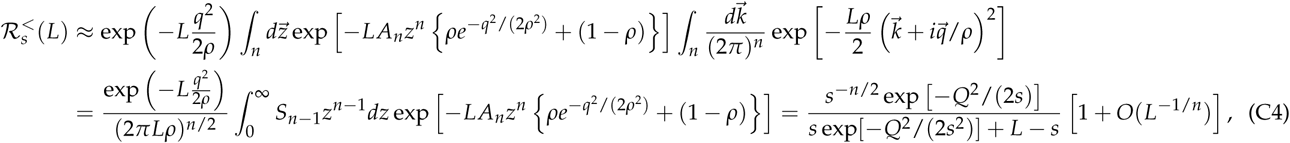
and the last step involves a change of variables *z* → *t* = S_*n*—1_*z^*n*^* /*n*. Since *L* appears in the integrand in the combination *Lz^n^*, the error that arises from neglecting terms of O(*z^n^*^+1^) is *L*^—1/^*^n^*. The leading order of Equation C4 was reported in Equation 34.

Now we move on to the calculation of 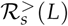, where the dominant contribution to 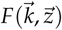 comes from a region where *z* ∼ O(*L*). Using 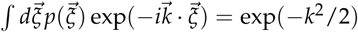 we calculate the integral 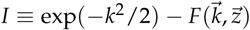 as

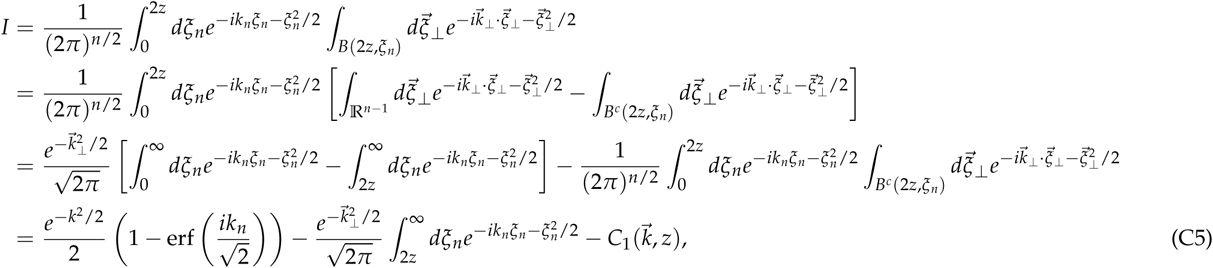
where we set 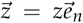 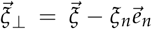 and 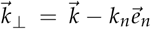 with 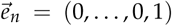, *B*(2*z, ζ_η_*) is an (*n* – 1) dimensional ball with radius 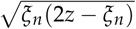 whose center is located at the origin, *B^c^* is the relative complement of *B* with respect to 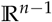, and 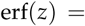 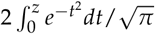is the error function. The definition of *C****_1_\*** is self-explanatory. Since

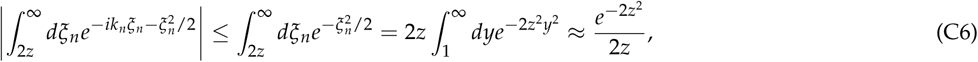
where we used the Laplace method for the asymptotic expansion, the leading finite *z* correction is expected to come from *C*_1_ for *n* > 1. Note that *C*_1_ is identically zero for *n* = 1. Thus we get

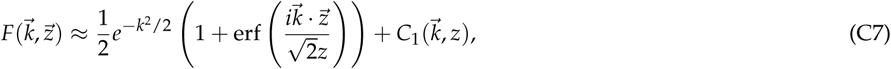
where *k_n_* is written as a projection of 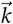 along the 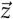 direction, 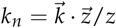 Since

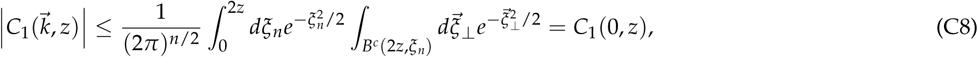
it is sufficient to find an approximate formula for C_1_(0,*z*) to determine the *z* dependence of 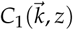. Using spherical coordinates in 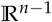, we get

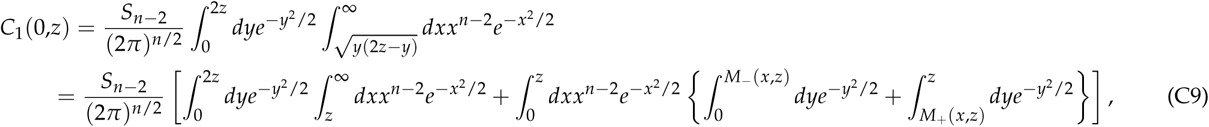
where 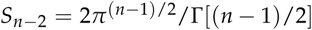 is the surface area of the unit (*n* - 2)-sphere. In the second term on the second line the order of integration was reversed and the integration boundaries 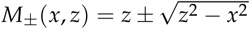 were introduced. Since the first integral 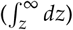and the third integral 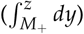 decrease exponentially with *z*, the main contribution to *C*_1_(0, *z*) comes from the second integral. Thus,

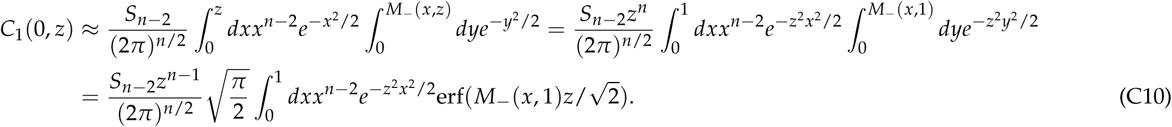

Since the last integral is dominated by the region *xz* **≤** 1, we can approximate M_—_(*x*,1)*z* ≈ *x*^2^*z*/2 ~ O(1/*z*) and erf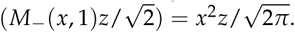 Finally, we get

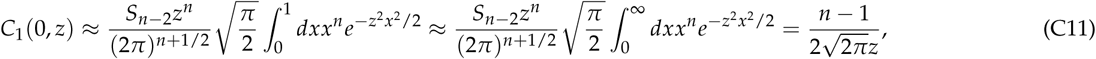
which also implies that 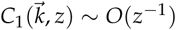. If we write

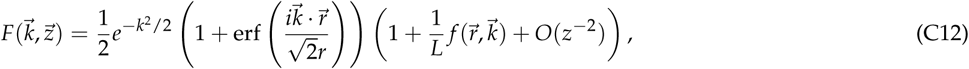
where 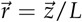 and *r* **=** *z****/****L*, then comparison with Equation C7 and Equation C11 shows that 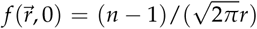 Inserting Equation C12 into Equation 32, it follows that

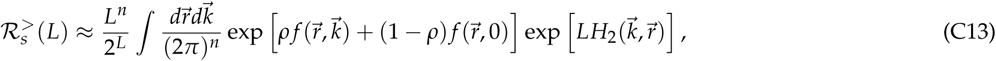

With

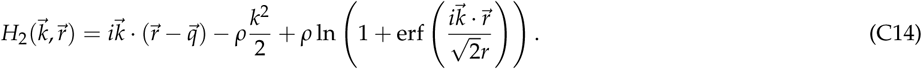

Now we employ the steepest descent method. For convenience, we set 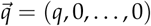. The saddle point satisfies the equations

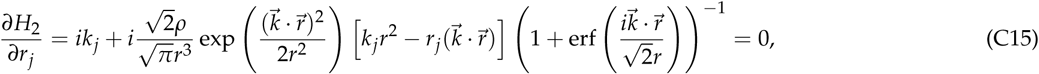

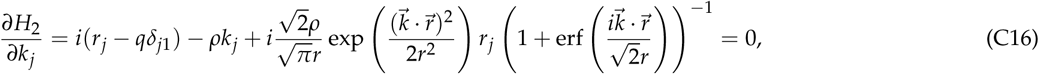
with the solution 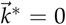 and 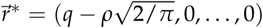. Note that there is no solution if 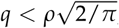 so the valid range of *ρ* has the upper boundary **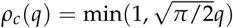.** The matrix of second derivatives around the saddle point 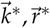 is

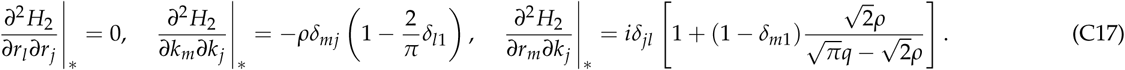

Thus, we get

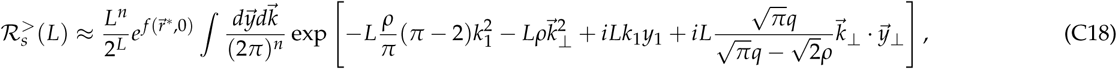
where 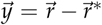,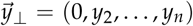, and 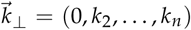. If we perform the integration over 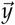 first, we obtain delta functions which make the integral over 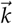 trivial. Finally, we arrive at

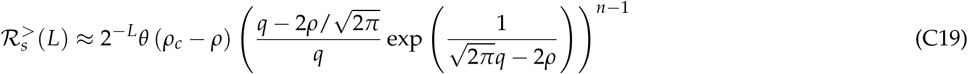
where *θ*(*x*) is the Heaviside step function defined by *θ*(*x* ≥ 0) = 1 and *θ*(*x* < 0) = 0.

To evaluate the corresponding contribution to the number of fitness maxima, 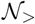, we replace the summation over s in Equation 33 by an integral over *ρ* = *s*/ *L* and use Stirling's formula to approximate the binomial coefficients. This yields

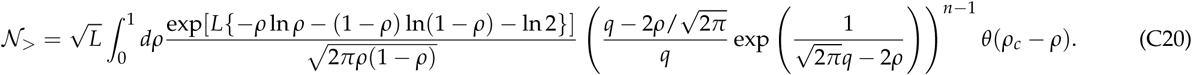

If 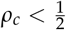 or 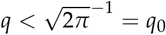the integral is dominated around *ρ* ≈ *ρ_c_*, which results in an exponential decrease with *L*. On the other hand, if 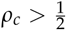 the integral is dominated around 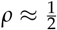 which gives

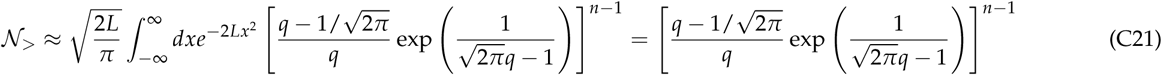
as reported in Equation 43.

## Appendix D: Derivation of Equation 47

In this appendix, we calculate the average number of fitness maxima 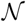 in the limit *n, L* → ∞ at fixed ratio *α* ≡ *n*/*L*. To this end, we write *I_τ_*, the probability for the genotype *τ* to be a local fitness maximum, using the Heaviside step function as

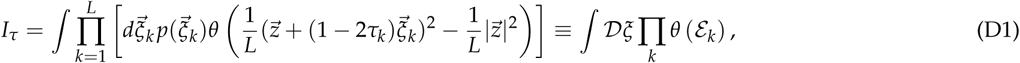
where 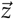 is determined by *τ* through Equation 3, 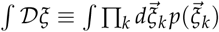 and ε*_k_* is defined as

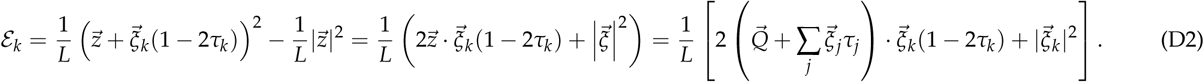

Note that the prefactor 1/*L* is introduced to make ε*_k_* finite in the limit *L* → ∞ and we have used that (1 – 2*τ_k_*)^2^ = 1. Applying the identity (Tanaka and Edwards 1980; Bray and Moore 1980)

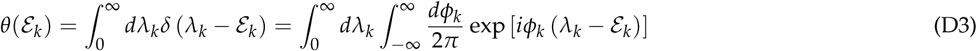
to Equation D1, the expected number of local fitness maxima reads

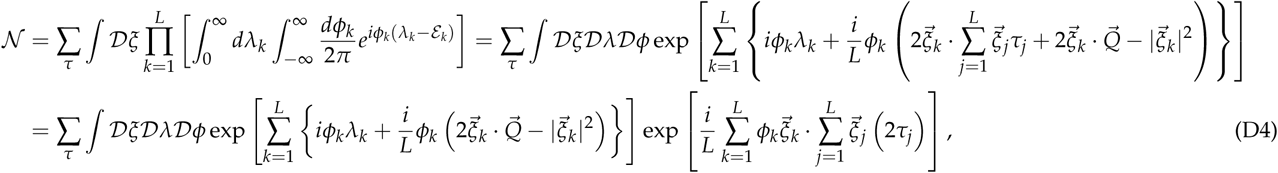
where 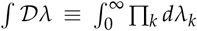, 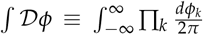 we made the change of variables 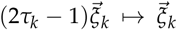 to arrive at the second equality. Using the identity

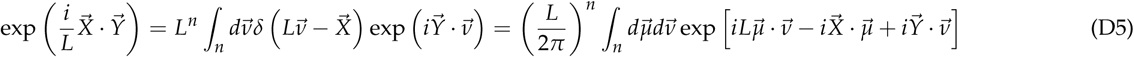
which is valid for any *n*-dimensional real vectors 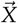 and 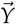, we can write the last term of Equation D4 as

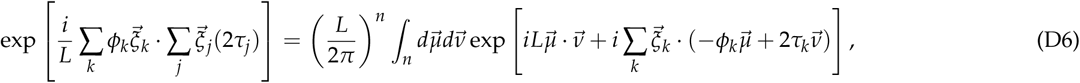
which gives

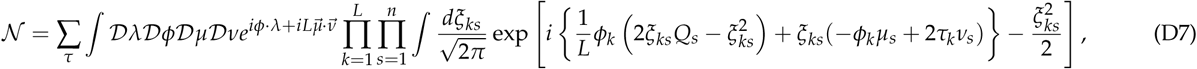
where 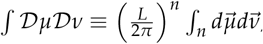, 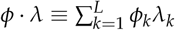 and ξ*_ks_, Q_s_, μ_s_, v_s_* are the *s*-th components of the vectors 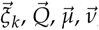 respectively.

Note that the integrals over the ξ*_k_*_s_'s become independent of each other. If we choose 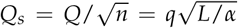 for all *s* and define *χ***=**2*Q_s_* /*L*, the integral over ξ*_ks_* becomes

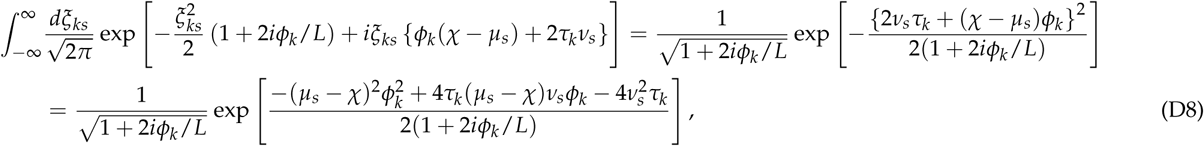
which, in turn, gives

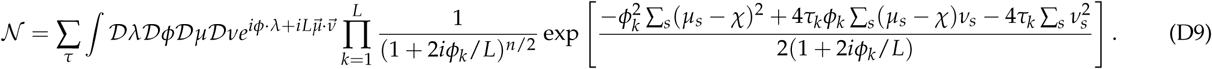

If we now insert the identity

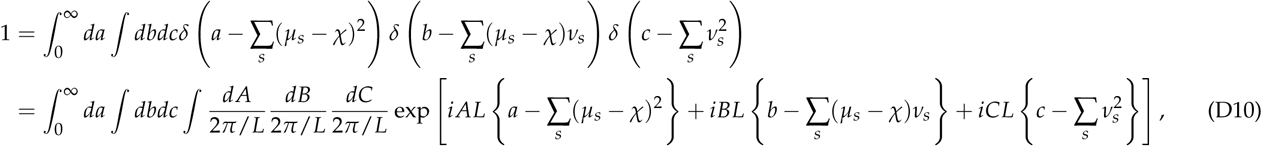
we can write

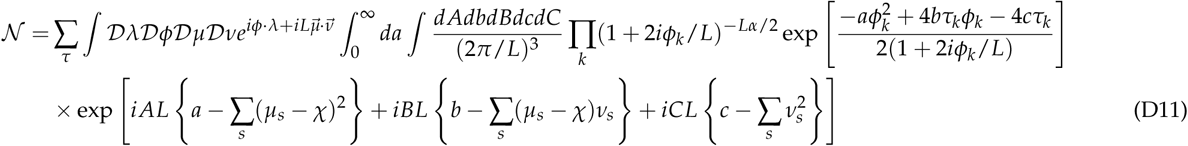
where we have replaced *n* by *Lα*. The integral domain of *a* is restricted to the positive real axis in order to ensure that the integral with respect to *ϕ_k_* in Equation D9 continues to be well-defined after the substitution. Performing the integrals over *μ_s_* and *v_s_*, we get

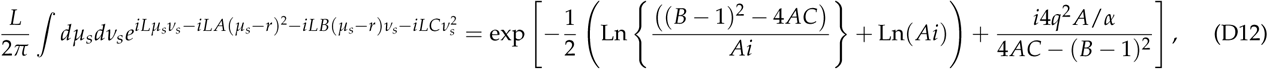
where Ln(*x*) is the principal value of the logarithm with argument in the interval ( –*π, π*] and the branch cut lies on the negative real axis.

Subsequently, the remaining integral over *ϕ_i_* and λ*_i_* can be readily evaluated as follows:

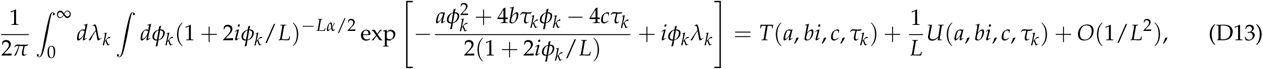

Where

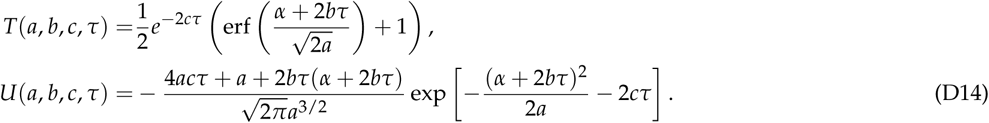

After summing over the τ*_k_*'s, we arrive at the equation

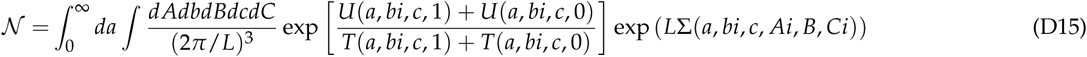

Where

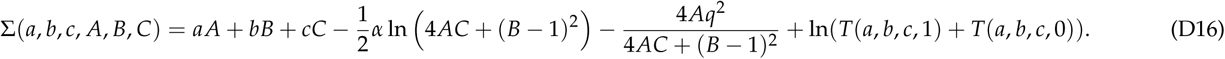

The remaining integrals are hard to evaluate analytically. Instead, we resort to the saddle point method to obtain an asymptotic expansion of the integral. Since Σ is the exponential growth factor of the number of local maximam which must be a real number, one expects that the saddle points of Equation D16 are formed for the real arguments of Σ. This suggests that we should make the changes of variables *b → b/i, A → A/i* and *C → C/i*. For large *L*, the integrals are then dominated by the saddle point (*a***\*,** *b***\*,** c**\*,** *A***\*,** *B***\*,** *C***\***) of Σ(*a, b, c*, A, B, C). If there is more than one saddle point, the one giving the largest value of Σ(*a, b, c*, A, B, C) has to be chosen. Then, the leading behavior of the number of maxima can be expressed in terms of the saddle point as

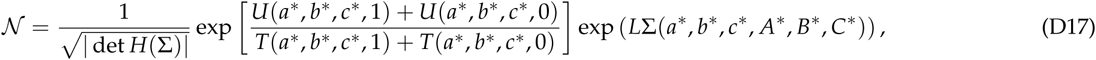
where Η(Σ) is the Hessian matrix around the saddle point. The reader may have noticed that the two principal values of the logarithm defined in Equation D12 are replaced by a real-valued logarithm in Equation D16, which can be dangerous in general. However, it can be shown that this substitution is indeed correct by verifying that (*B*\* – 1)^2^ + 4*A*\* *C*\* is always positive for all saddle points of Equation D16, and thus the imaginary arguments always cancel each other out.

Now, let us evaluate the saddle point conditions. The derivatives of Σ with respect to *A*, *B*, *C* are

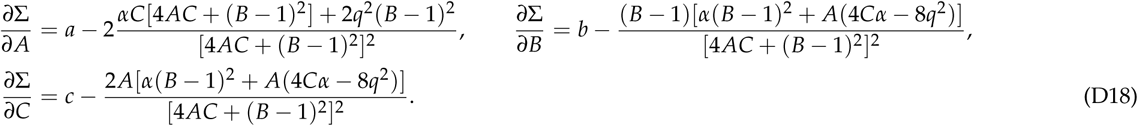

By requiring that the above three equations are zero at the saddle point, we get

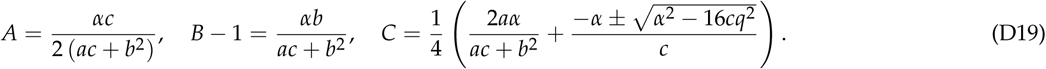

The two solutions of *C* force us to perform a two-fold analysis for the remaining integrals since we cannot *a priori* determine which solution will yield the correct saddle point. Instead, we introduce another real number 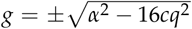 which is allowed to take both signs. Then, by imposing the functional relation 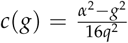, both solutions are covered by a single analysis. In this way, the saddle point is obtained in terms of *g* instead of *c*. Finally, substituting this solution into Equation D17 gives Equation 47.

## Appendix E: Mean phenotypic distance *z* in the joint limit

In this appendix, we will associate the fixed point value *a*^*^ of the variable *a* entering the complexity function Equation 47 with the mean phenotypic distance *z*\*. To this end, we first consider the probability density *Ρ*(τ, *a*) that a genotype *τ* whose phenotypic vector is of squared magnitude *L*^2^*a*/4 is a local maximum. Formally, we can write

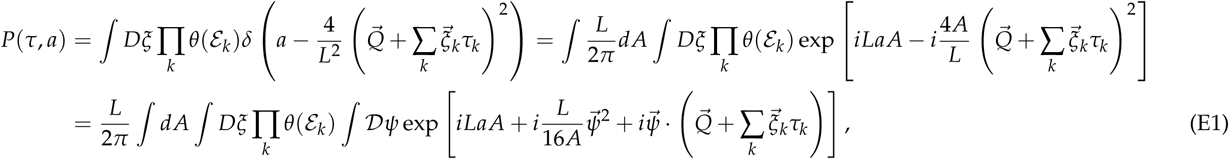
where we have used the identity 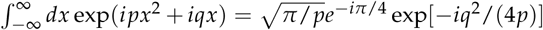 for *p* > 0,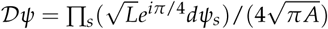 and the notation is the same as in Appendix D. Following the same procedure in the previous appendix, we get

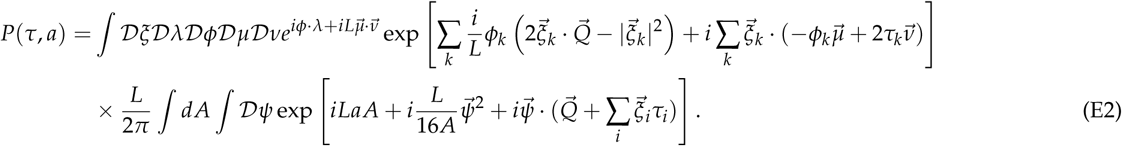

By shifting 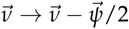 and integrating over 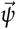, we have

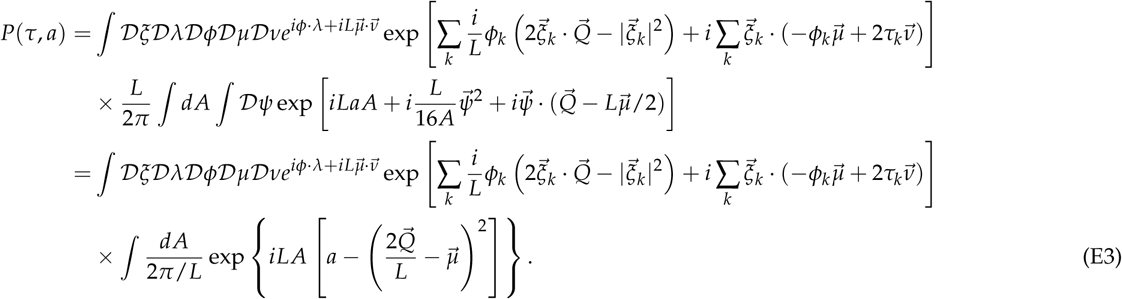

Since we have set 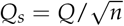 for *s* = 1˙,*n* last integral becomes

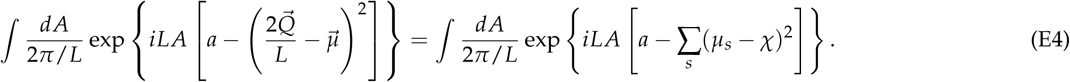

Since 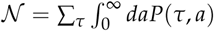, by applying the manipulations of Appendix D to Equation E3 we arrive at the same integral form as in Equation D10. Since Σ_τ_ *P*(τ, *a*) is the mean number of local maxima whose phenotypic vectors have squared magnitude *a*, we see that the saddle point *a*\* of Equation D16 determines the mean phenotypic distance *z*\* through

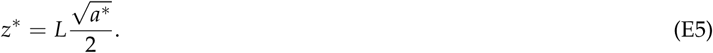

This shows in particular that *z*^*^ is linear in *L*.

## Appendix F: Mean genotypic distance *ρ\** in the joint limit

To have access to the information about the typical value of the genotypic (Hamming) distance of a local fitness maximum from the wild type, we rewrite Equation D15 as

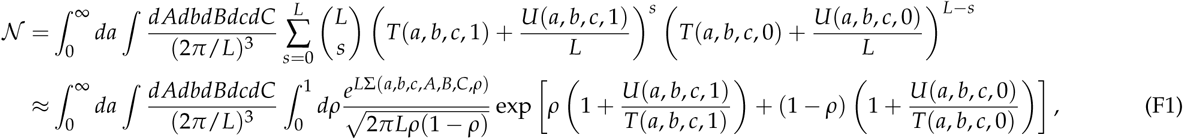
where we have rearranged the summation Σ_τ_ as 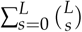 taking advantage of the inherent permutation symmetry, Stirling's formula has been used to evaluate the binomial coefficients, Σ_s_ is approximated as 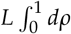 with *s* = *Lρ*, and

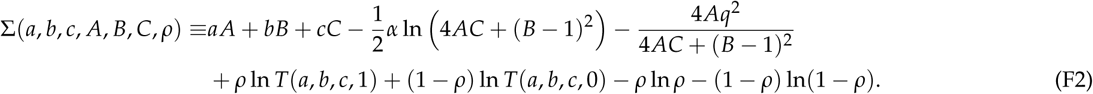

The saddle point equations for this expression involve seven variables including *ρ*. Since the saddle point equations for *A, B, C* are the same as Equation D18, we may again insert Equation D19 into Equation F2, which yields

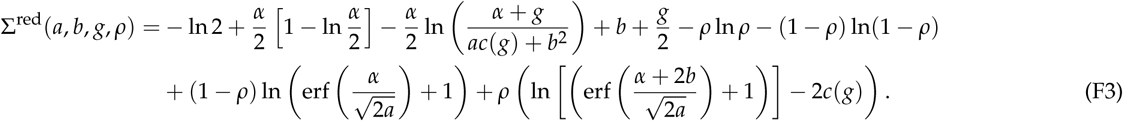

Since

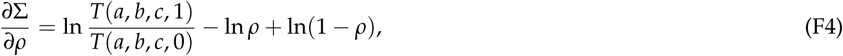
the saddle point value of *ρ*^*^ is

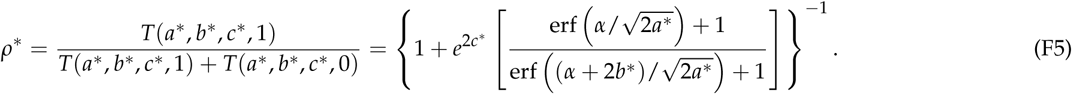

By inserting *ρ*\* into the saddle point equations for *a, b, c*, one can easily see that the final equations are the same as those derived from Equation D16.

## Appendix G: Derivation of Equation 50

The determination of the solution describing regime III relies on the intuition that as *α* becomes large, the fitness landscape is asymptotically linear with the wild type being the global fitness maximum, as demonstrated in **Sign epistasis** for *L* = 2. This suggests an ansatz where *a*\* is close to 4*q*^2^, which corresponds to the wild type phenotypic distance as shown in Equation E5. Given this clue, one can additionally find that

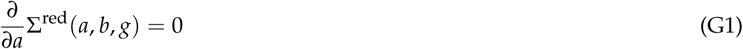
is solved by *a* = 4*q*^2^, *b* = –*α* and *g* = *α*. Furthermore, if we evaluate the remaining saddle point conditions around this point, we find that this solution fails to solve them by a slight margin,

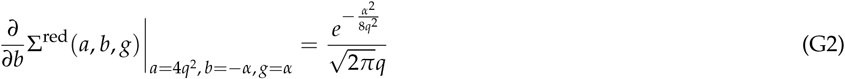

And

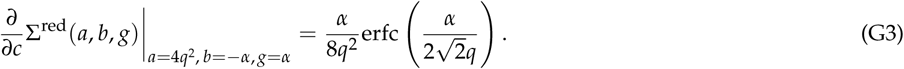

Given the fact that 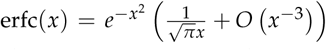, these non-vanishing terms are seen to be of the order of 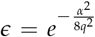. Hence, it is sufficient to consider an expansion around the zeroth order solution of the form Σ^red^ (4*q*^2^ + *A*_1_*∈*, –*α* + *A*_2_*∈, α* + *A*_3_*∈*) to show that Equation 50 satisfies the saddle point conditions Equation 48. To this end, we first focus on the derivatives with respect to *A*_1_ and *A*_2_,

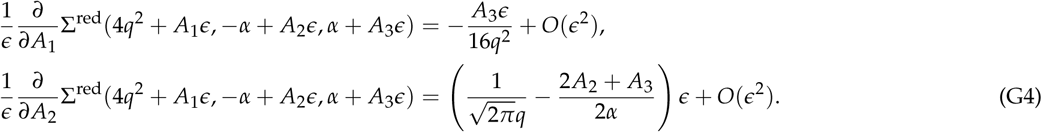

The vanishing contributions in *∈* imply that the zeroth order solution (4*q*^2^, –α, α) satisfies the first two saddle point conditions. Additionally, we find that the corrections of the order O(*∈*) are *A*_3_ = 0 and 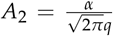 Since *A*_3_**=** 0 to leading order, the saddle point equation with respect to *g* should be evaluated to order O(*∈ ^2^*). This yields

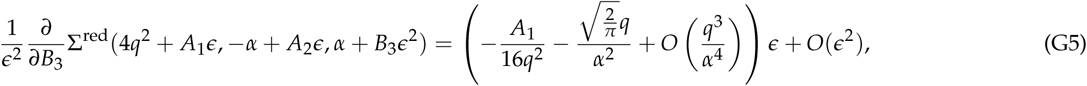
and subsequently, *A*_1_ is solved to be 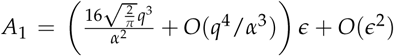 Finally, by inserting the solutions *A*_1_, *A*_2_ and *A*_3_ as well as the zeroth order solutions into Equation F5, the solution for *ρ*^*^ is found to be

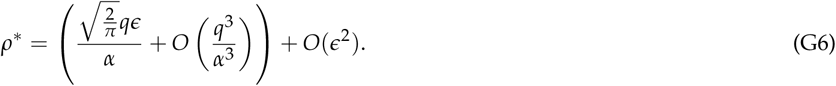

